# A cross-species analysis of neuroanatomical covariance sex differences in humans and mice

**DOI:** 10.1101/2024.11.05.622111

**Authors:** Linh Pham, Elisa Guma, Jacob Ellegood, Jason P. Lerch, Armin Raznahan

## Abstract

Structural covariance in brain anatomy is thought to reflect inter-regional sharing of developmental influences - although this hypothesis has proved hard to causally test. Here, we use neuroimaging in humans and mice to study sex-differences in anatomical covariance - asking if regions that have developed shared sex differences in volume also show shared sex differences in volume covariance. This study design illuminates both the biology of sex-differences and theoretical models for anatomical covariance – benefitting from tests of inter-species convergence. We find that volumetric structural covariance is stronger in adult females compared to adult males for both wild-type mice and healthy human subjects: 98% of all comparisons with statistically significant covariance sex differences in mice are female-biased, while 76% of all such comparisons are female-biased in humans (q < 0.05). In both species, a region’s covariance and volumetric sex-biases have weak inverse relationships to each other: volumetrically male-biased regions contain more female-biased covariations, while volumetrically female-biased regions have more male-biased covariations (mice: r = -0.185, p = 0.002; humans: r = -0.189, p = 0.001). Our results identify a conserved tendency for females to show stronger neuroanatomical covariance than males, evident across species, which suggests that stronger structural covariance in females could be an evolutionarily conserved feature that is partially related to volumetric alterations through sex.

**SIGNIFICANCE STATEMENT:** Structural covariance is a potent readout of coordinated brain development, but hard to probe experimentally. We use sex differences as a naturally occurring test for developmental theories of structural covariance – adopting a cross-species approach for validation and translational benefit. Brain MRI reveals two conserved features of anatomical covariance across humans and mice: (i) tighter inter-regional coordination of brain development in females as evidenced by stronger volume covariance; (ii) a tendency for female-biased covariance to involve regions that are smaller in females – suggesting a previously unknown counterbalancing between these two distinct modes of sex-biased brain organization. These findings advance understanding of coordinated brain development and sex difference in a cross-species framework – facilitating future translational research on both topics.

## INTRODUCTION

Structural covariance refers to the phenomenon in which variable biological structures in a population scale together across individuals. The degree of covariation between two structures is typically taken as evidence for how strongly they relate at some unmeasured levels of biology (Galton, 1888). It is thought that observed anatomical covariance between regions in adulthood reflect their shared developmental influences in earlier life. This idea is based on published observations that higher inter-regional structural covariance tracks with stronger inter-regional coupling in inter-regional connectivity (Andrews et al., 1997; Gong et al., 2012), gene co-expression (Romero-Garcia et al., 2018; Yee et al., 2018), and developmental tempos (Raznahan et al., 2011; Alexander-Bloch et al., 2013). A direct way to test this hypothesis would be to experimentally manipulate the developmental influences of brain regions sets across multiple individuals and then estimating structural covariance across the group at a later time point. However, this experimental approach is technically challenging and could introduce unintended changes that confound interpretations of developmental effects on covariance formation.

Sex differences in brain organization serve as an alternative approach for testing the “shared influences” model of structural covariance. Experimental data have identified rodent brain regions with reproducible volumetric sex differences (Corre et al., 2016) through regional action of male-specific hormonal effects (Gorski et al., 1978; Segovia et al., 1984; Roos et al., 1988; Hines et al., 1992; Williams et al., 2001; Forger et al., 2004; Morris et al., 2008). These volumetric sex differences include male-biased volume in the bed nucleus of the stria terminalis (BNST), olfactory bulb, medial amygdala, and female-biased volume in the anteroventral periventricular nucleus (AVPV) (Segovia et al., 1984; Roos et al., 1988; Hines et al., 1992; Williams et al., 2001; Forger et al., 2004; Morris et al., 2008). Humans also show highly reproducible sex differences in regional brain anatomy (Liu et al., 2020; DeCasien et al., 2022) that presumably reflect sex-biased regional brain development. Some of these biases share directionalities with mice, such as the male-biased BNST and medial amygdala (Guma et al., 2024). If adult neuroanatomical covariances arise through an inter-regional sharing of developmental influences, then one would expect that volume covariance tends to be more sex-biased amongst brain regions that are sex-biased in mean volume than amongst those that are not. Thus, the study of sex-differences in neuroanatomical covariance not only sheds light on an understudied axis of sex-biased brain organization but also provides a naturally occurring probe for developmental models of covariance. To date, however, there is pronounced heterogeneity in results across those few studies that have tested for sex-biased structural covariance in humans (Mechelli et al., 2005; Persson et al., 2014; Wierenga et al., 2018; Seitz et al., 2019; Ge et al., 2021; Vijayakumar et al., 2021; Shi et al., 2023) without comparisons to sex-differences in regional volume. Recent works in larger cohorts suggest cortical thickness covariance patterns are stronger in males while volume covariance patterns are stronger in females (Wierenga et al., 2022; Yang et al., 2023). To our knowledge, there are no published studies explicitly examining normative sex-biased neuroanatomical covariance in mice.

Here, we use cross-species structural magnetic resonance imaging (sMRI) to map sex-biased brain volume covariance and sex-biased volume. We test the hypothesis that these features are non-randomly correlated across the brain. Addressing this question requires computing sex differences in covariance between all pairs of regions within each species – which enables us to address several related questions including: (i) whether there is a tendency towards stronger volume covariance in one sex within each species, (ii) whether there is an inter-species differences in the strength of inter-regional neuroanatomical covariance, (iii) what specific brain regions and inter-regional pairs show sex-biased covariance in each species. Taken together, our work provides the first comparative analysis of sex-biased neuroanatomical covariance in humans and mice. Our results inform dominant developmental theories for the emergence of structural covariance and expand our comparative understanding of sex-biased mammalian brain organization.

## METHODS

### Acquisition and processing of murine neuroimaging data

Our study includes structural MRI (sMRI) brain scans from 423 mice acquired at the Mouse Imaging Centre in Toronto. Scans were performed on the same 7T multichannel scanner with either an insert gradient (6 cm inner bore diameter magnet) or an outer gradient (30 cm diameter bore diameter magnet) (Agilent Inc., Palo Alto, CA). Mice were transcardially perfused using a standard protocol across all cohorts (Spring et al., 2007; Lerch et al., 2011; Cahill et al., 2012). Brains were kept in the skull and fixed to avoid distortions during imaging. All animal procedures were approved by the ethics committees of their originating labs and the animal care committee at The Centre for Phenogenomics (AUP-0260H) at the University of Toronto.

As the mouse data for this study was collected over 10+ years, the MRI pulse sequences were optimized over that period to increase the scanning throughput to enable 16 mice to be scanned in one session, and/or to improve resolution and increase gray/white matter contrast in each scan (Lerch et al., 2011; Ellegood et al., 2015). The following three MRI sequences were used in overnight scans throughout the studies included here. (1) 3 brains scanned in parallel per session -- T2-weighted fast spin echo (FSE): TR = 325 ms, TE = 10 ms/echo for 6 echoes. The center of k-space is acquired on the 4th echo. Field-of-view (FOV) = 14 x 14 x 25 mm^3^. Matrix size = 432 x 432 x 780. Image resolution = 32 µm isotropic voxels. (2) 16 brains scanned in parallel per session (sequence 1) -- T2-weighted 3D FSE: TR = 2000 ms, echo train length = 6, TE_eff_ = 42 ms. FOV = 25 x 28 x 14 mm^3^, matrix size = 450 x 504 x 250. Image resolution: 56 µm isotropic voxels. Oversampling in the phase encoding direction by a factor of 2 was applied to move ghosting artifacts from k-space discontinuity to FOV edges. FOV was cropped to 14 mm after image reconstruction. (3) 16 brains scanned in parallel per session (sequence 2) -- T2-weighted 3D FSE: TR = 350 ms, TE = 12 ms/echo for 6 echoes. Cylindrical 3D k-space acquisition. FOV = 20 x 20 x 25 mm^3^, matrix size = 504 x 504 x 630. Image resolution: 40 µmm isotropic voxels (Spencer Noakes et al., 2017). Of note, these sequences were evenly distributed between male and female mice.

Structural MRIs were registered and warped to an average study mouse template using deformation-based morphometry (Collins et al., 1994; Avants et al., 2009; Avants et al., 2011; Eskildsen et al., 2012; Friedel et al., 2014). Log-transformed Jacobian determinants for each voxel were calculated and used to determine voxel volume differences between individual mouse brains with the averaged brain (Chung et al., 2001). ROI volumes were calculated as the sum of volume differences for each voxel within the ROI. This process used the MAGeT algorithm (Chakravarty et al., 2013; Pipitone et al., 2014) and resulted in 336 unique brain regions from previously published atlases (Dorr et al., 2008; Richards et al., 2011; Ullmann et al., 2013; Steadman et al., 2014; Qiu et al., 2018). 255 of these regions were grey matter and included in the study.

The imaged mice consist of C57BL6J (n = 152) and C57BL6N (n = 271) wild-type controls from separate studies that each compared wild-type controls with mutations of a different autism-related risk gene (Ellegood et al., 2015). For the purposes of this study, we only included wild-type cohorts which had at least 5 male and 5 female mice surviving a quality assessment procedure to flag and remove outliers. This procedure involved an initial visual quality control was performed to ensure accurate registration and segmentation, followed by an outlier detection process.

Regional volumetric measures for the full set of 432 sMRI scans was then subjected to batch control using ComBat (*sva* library in R) to correct for variability between strain and ASD gene cohort of origin (Johnson et al., 2007; Fortin et al., 2017; Fortin et al., 2018; Leek et al., 2022). Final animal characteristics are detailed in Table 1.

**Table 1.**
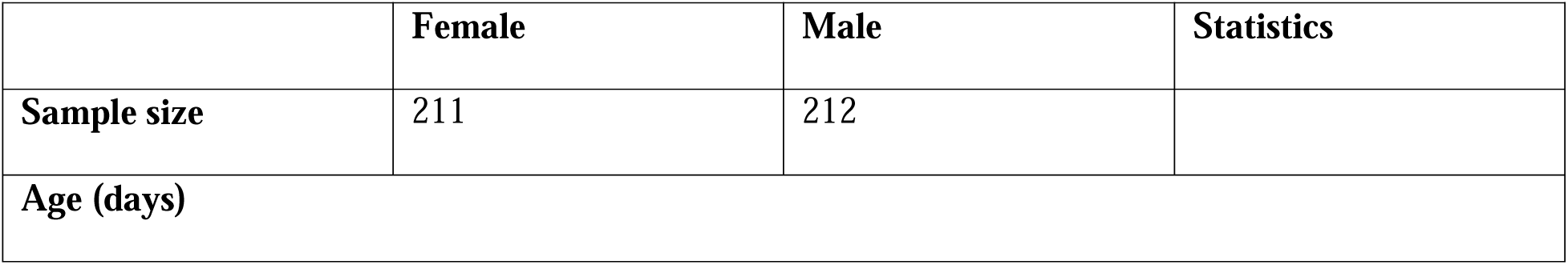
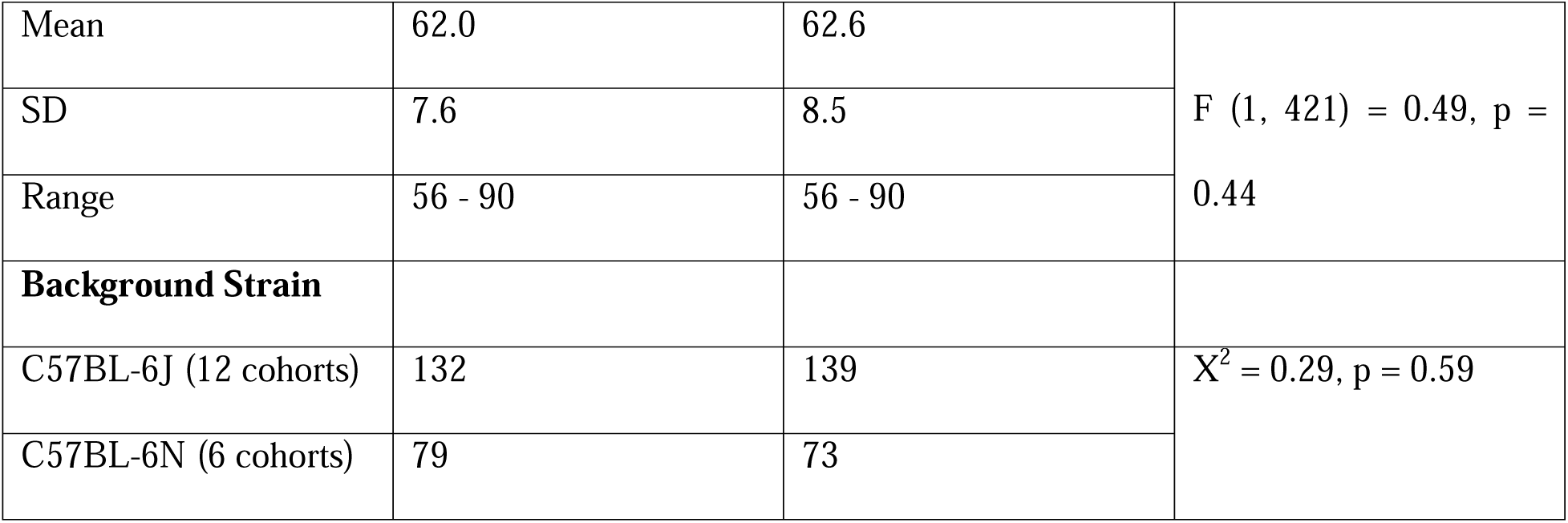
Demographics for mouse sample.

### Acquisition and processing of human neuroimaging data

This study includes 436 human sMRI brain scans from the Human Connectome Project 1200 release. Scans were obtained using an MR750 3-T (General Electric) whole-body scanner (MP-RAGE-T1: TE 2.14 ms, TR 2400 ms, flip angle = 8°, FOV 224 × 224 mm2, scan time = 7:40 min, voxel size = 0.7 mm isotropic) with a 32-channel head coil (176 continuous sagittal slices with 256 x 256 in-plane matrix and 1 mm slice thickness). Additional recruitment procedures and acquisition parameters are detailed in the original publication (Van Essen et al., 2012; Glasser et al., 2013). Information on how to obtain HCP data can be found here (https://www.humanconnectome.org/study/hcp-young-adult/document/wu-minn-hcp-consortium-restricted-data-use-terms). 1110 unique subject scans were visually inspected and removed if obvious registration and/or segmentation issues were detected. Euler numbers – indicators of brain topological reconstruction quality – were also measured forx each scan using the image preprocessing steps described below (Dale et al., 1999). Scans with FreeSurfer-estimated Euler numbers less than -217 were excluded from further analyses (Rosen et al., 2018). From the remaining 1030 subject scans, we randomly selected one person per family, based on distinct mother ID and father ID, to yield 436 unique and unrelated subjects.

All remaining subjects underwent an outlier flagging process using maximal Cook’s distance. Briefly, pairwise linear models were created between all brain structures using mixed sex data. Subjects’ maximum Cook’s distances from all linear models are recorded. This served to identify whether a subject appeared to be unusually influential on the relationship between any pairwise correlations. A subject was deemed to have an unusually influential effect on pairwise relationships if its maximum Cook’s distance was greater than the standard cut off: 3 z-scores of the maximum Cook’s distance for all subjects. Such subject’s scan would be flagged for a secondary visual review. If a further review did not show abnormalities in the scan, it was kept in the data set. Final participant characteristics are detailed in Table 2.

**Table 2.**
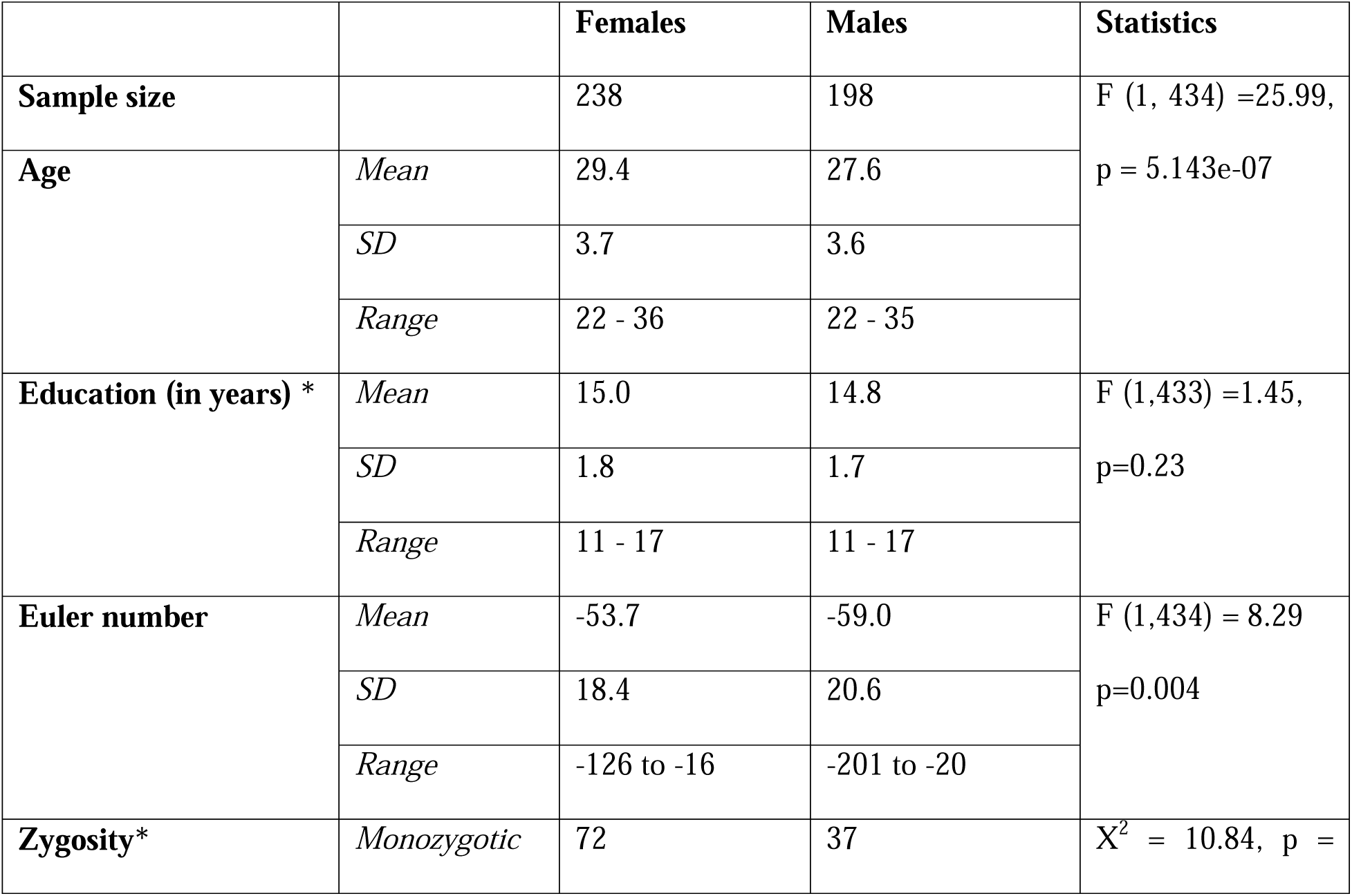
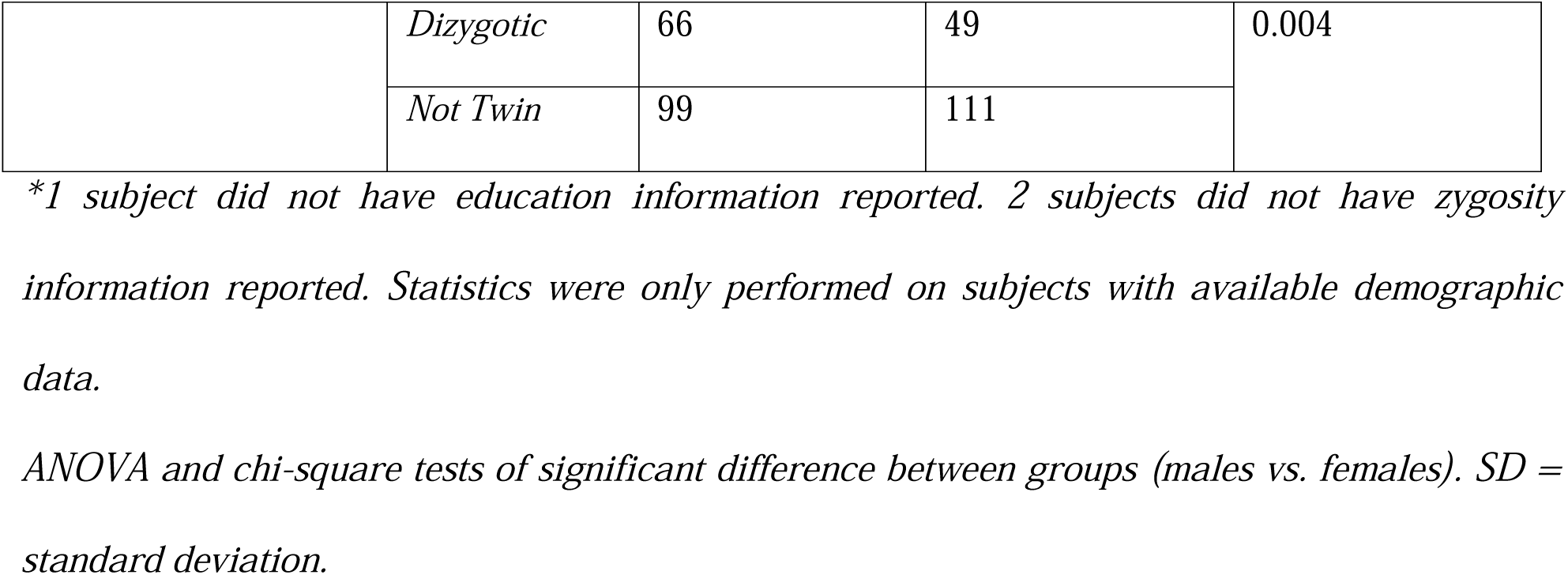
Demographics for human sample.

The PreFreesurfer pipeline was used to preprocess T1-weighted structural MRI data (Glasser et al., 2013). Freesurfer 7.1.0’s (Fischl, 2012) *recon-all* and *highres* commands were used to reconstruct and parcellate the cortex at the original data resolution (Dale and Sereno, 1993; Fischl et al., 1998; Dale et al., 1999; Fischl et al., 1999a; Fischl et al., 1999b; Fischl and Dale, 2000; Fischl et al., 2001; Fischl et al., 2002; Kuperberg et al., 2003; Fischl et al., 2004a; Desikan et al., 2006; Han et al., 2006; Jovicich et al., 2006; Reuter et al., 2010; Zaretskaya et al., 2018). The pipeline can be downloaded here (http://surfer.nmr.mgh.harvard.edu/). Cortical volumes were extracted using the *mri_anatomical_stats* utility. 360 regions from the Glasser Human Connectome Project were generated using this procedure (Glasser et al., 2016). 359 regions from this atlas were used for subsequent analyses.

Subcortical and hippocampal segmentation was performed by first assigning one of 39 labels from the FreeSurfer ‘aseg’ feature to each voxel (Fischl et al., 2002; Fischl et al., 2004b). 19 of these labels were gray matter structures and were included in subsequent analyses. Additional segmentations of sex-biased nuclei in the hippocampal subfield, amygdala sub-nuclei, and brainstem were made using FreeSurfer joint segmentation of these subfields (Iglesias et al., 2015b; Iglesias et al., 2015a; Saygin et al., 2017). Segmentations for classically sex-biased BNST and hypothalamic nuclei were made under a different atlas that is not available within FreeSurfer (Neudorfer et al., 2020) (https://zenodo.org/record/3942115). Hypothalamic atlas labels were registered to the study’s average template, and deformation-based morphometry was applied to warp each subject’s image to the study’s template (Devenyi, 2024) (https://github.com/CoBrALab/optimized_antsMultivariateTemplateConstruction). The Jacobian determinant from this process was used to calculate the volume change from the template at each voxel in the region of interest (ROI). A summation of these changes within the ROI results in its volume measurement.

### Comparing regional volume covariance between males and females in each species

To compare region of interest (ROI) covariance across sex in each species, we split the data for each species by sex and regressed age out of ROI volumes for both species using the following model:

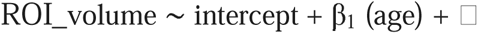

Residuals from this model were used to compute all pairwise inter-regional volume correlations within males and females. The means of these correlations across all pairwise relationships were directly compared between males and females using t-tests and reported 95% confidence intervals (CI). We then subtracted the male correlation matrix from the female to derive a measure of sex differences in correlation for all region pairs (with positive values indicating a larger correlation in males vs. females). The statistical significance of sex differences in correlation for each unique pair of regions was determined by repeating the above process 1000 times with sex being permuted across individuals for each iteration. This procedure yielded a vector of 1000 null values for each pairwise correlation sex differences and we derived empirical p-values for these observed sex differences against these nulls. Empirical p-values were corrected for multiple comparisons across edges using the False Discovery Rate (FDR) (Benjamini and Hochberg, 1995; Benjamini and Yekutieli, 2001) correction with q (the expected proportion of falsely rejected nulls) being set at 0.05.

### Examining the relationship between sex differences in volume covariance and sex differences in regional volume

Sex differences in regional volume were estimated as follows: ROI and total grey matter tissue volumes (TTV) were z-scored across individuals, then input into the following models to estimate the effect of sex on the mean volume of each brain region (given by the β_1_ coefficient in the model below):

Mice: ROI_volume ∼ intercept + β_1_(Sex: male vs female) + β_2_(age) + β_3_(TTV) + β_4_(Background Strain) + □

Human: ROI_volume ∼ intercept + β_1_(Sex: male vs female) + β_2_(age) + β_3_(TTV) + β_4_(Euler number) + □

Positive beta coefficients in the model indicate male-biased regional volumes. p-values associated with the β_1_ coefficients from each of these models were corrected for multiple comparisons across the number of brain regions in each species using FDR with q < 0.05.

The relationships between covariance and volumetric sex differences were assessed using several complimentary approaches. First, we selected three classically sex-biased regions in mice – the BNST, medial amygdala, and olfactory bulb – and asked if there are any covariance sex differences among these pairings. Statistical significance was calculated using the previously described permutation pipeline and Bonferroni corrected for the total number of comparisons made within this analysis (p = 0.05). Second, we used the full correlation sex differences matrix in each species to estimate the mean sex differences in volume correlation per brain region (averaging the sex differences in its correlation with all other regions) - once using all pairwise correlation sex differences, and again using just those pairs deemed statistically significant in covariance sex differences through permutation testing. We examined the distribution of these properties across the brain of each species and noted those brain regions showing both sex differences in mean volume and sex differences in volume covariance. Third, we used network visualization to specify sets of brain regions showing prominent sex differences in anatomical covariance. Specifically, we identified all region pairs with statistically significant covariance sex differences, converted these pairings into a graph with nodes (regions) and edges (covariance sex differences), and visualized the largest connected components of these graphs in each species to determine their contents and any included nodes that also show sex-biased volume. To capture a similar number of nodes across species for these graphical representations we examined the two largest connected components in mice and the single most connected component in humans. Fourth, we tested if inter-regional variation in the mean sex difference in volume covariance per brain region was correlated with inter-regional variation in the effect size of volumetric sex differences (β_1_coefficients). These correlations were run twice in each species – once using the regional means for the absolute sex differences in correlations and once using regional means for signed sex differences in correlations. The absolute value test asks if regions with a larger volumetric sex bias tend to show larger sex-differences in their inter-correlations - irrespective of the “direction” (i.e. male-biased vs, female-biased) of these differences. In contrast, the signed test asks if regions with a more strongly biased volume in one sex versus the other tend to show similarly signed sex differences in their covariance (i.e. regions with male-biased in volume tend to show male-biased in their covariance). The statistical significance of these correlations between regions’ sex differences in covariance and regional sex differences in volume was assessed by comparing observed correlations with a distribution of 1000 null correlations from permutations of sex within species (performed on the NIH HPC Biowulf cluster -- http://hpc.nih.gov). These global correlations between sex-biased regional volume covariance and sex-biased regional volume represent the core brain-wide test of our primary motivating hypotheses based on developmental theories for structural covariance: that brain regions exposed to sex biased influences on their volume in development (manifest as sex-biased mean volume in adulthood) would therefore be expected to show more prominent sex-biased covariance than brain regions lacking a sex difference in volume.

### R versions and packages

All analyses presented in this paper were performed using R version 4.2.3 unless Biowulf handled the computation, in which case R version 4.2.1 was used (R Core Team, 2023). Packages used for all analyses can be found in the references section (Lerch et al., 2017; cocoframer, 2018; Lander, 2018; Mowinckel and Vidal-Piñeiro, 2019; Wickham et al., 2019; Glur, 2020; Kuhn and Wickham, 2020; Wickham, 2020; Landau, 2021; Mowinckel and Vidal-Piñeiro, 2022; Lerch, 2023; Mowinckel and Vidal-Piñeiro, 2023; Robinson et al., 2023; Pedersen, 2024). Data cleaning and analyses codes can be found at github.com/phamlk/cross-species-covariance-sex-differences.

## RESULTS

### Mean structural covariance is slightly stronger in females than males for both mice and humans

For both species, we first compared the means of all correlations in each sex prior to identifying specific interregional pairs with large covariance sex differences. Our comparison showed that mean interregional covariance is stronger in females than males, with a small magnitude for the mean between sex difference (Δ) in covariance (**Figure. 1 A - B**, Δ mice: 0.043, 95% CI: 0.041 - 0.045; Δ humans: 0.016, 95% CI: 0.015 - 0.017). After correction for multiple comparisons across region pairs, we identified 44 pairs of regions with statistically significant sex-biased covariance in mice (0.14 % of all pairwise relationships in the mouse brain; 43 stronger covariance in females) and 100 such pairs in humans (0.10 % of all pairwise relationships in the human brain: 71 stronger covariance in females). As expected, the mean of covariance strengths for these pairs differed between the sexes for both species - with a larger magnitude than was seen when considering all pairs (**Figure 1 C - F**, Δ mice: 0.293, 95% CI: 0.235 - 0.353; Δ humans: 0.156, 95% CI: 0.108 - 0.204). Of note, the average within-sex correlation was consistently higher in mice than in humans (all comparisons: 0.275 in male mice, 0.196 in male humans. 0.318 in female mice, 0.212 in female humans; significant comparisons: 0.129 in male mice, 0.106 in male humans. 0.422 in female mice, 0.261 in female humans). The full within-sex structural covariance matrices and lists of pairwise covariance sex differences can be found for each species in **Extended Figures 1-1, 1-2 and Extended Tables 1-1, 1-2.** As part of these analyses, we also tested the expectation that sex differences in volume correlation are more pronounced in region pairs showing weaker within sex volume correlation (as simulated in **Extended Figure 1-3**). We confirmed the presence of an inverse relationship between covariance strength and covariance sex differences when considering pairings at various correlation sex differences significance thresholds. The results of these analyses are provided in **Extended Figure 1-4**. Taken together these results indicate that regional volume covariance is stronger in mice than humans for both sexes, and that within each species, females tend to show stronger structural covariance than males. For both species, these sex-differences in structural covariance are statistically significant for a small subset (<0.15%) of all possible inter-regional pairings in each species, with the largest sex differences occurring between those inter-regional pairings that show weaker structural covariance in each sex.

**Figure 1.**
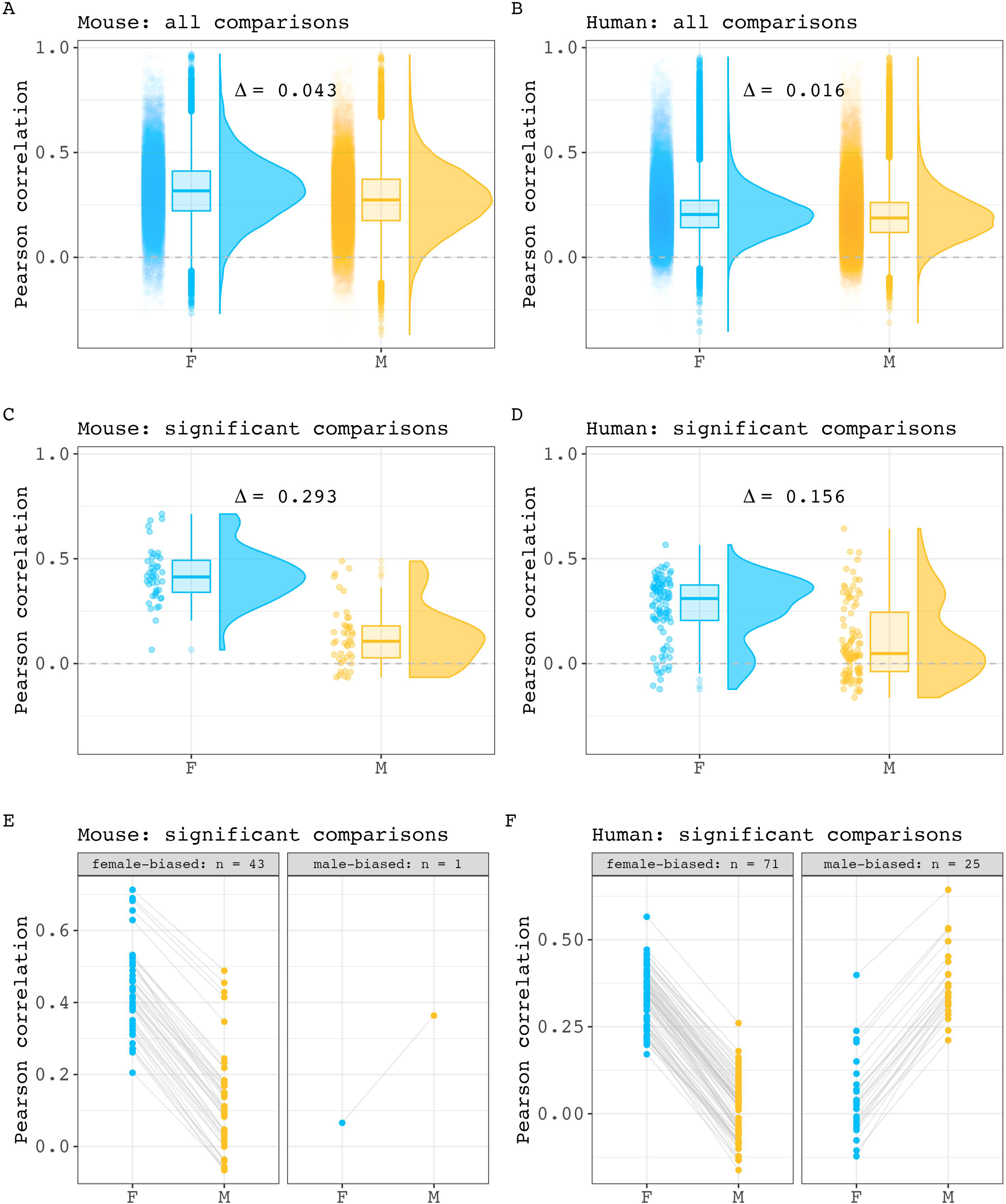
Inter-regional volume correlation distributions across sex in mice and humans. **A, B)** Comparison of all within sex correlation values for mouse (A) and human (B) (Δ mouse = 0.043, 95% CI: 0.041 - 0.045. Δ human = 0.016, 95% CI: 0.015 - 0.017). Each point represents a pairwise correlation value. Box plots and density plots are shown for distribution visualizations. **C, D)** Comparison of within-sex correlation values for region pairings with statistically significant covariance sex differences in mouse (C) and human (D) (Δ mouse = 0.293, 95% CI: 0.235 - 0.353. Δ human = 0.156, 95% CI: 0.108 - 0.204). **E, F)** Pairwise visualization of comparisons with significant sex differences in mouse (E) and human (F). Each point represents a pairwise correlation in either male or female. A connecting line between two points are shown to connect a pairwise correlation value in one sex and with its equivalent pairing in the other sex.

### Cross-brain analysis reveals a weak association between interregional sex differences in volume covariance and sex differences in regional volume

We took several complementary approaches to examining the relationship between regional sex differences in brain volume covariance and regional sex differences in brain volume. First, we selected 3 classical regions with the largest and best-replicated sex differences in volume in mice – BNST, olfactory bulb, medial amygdala – and examined sex differences in covariance between these structures. None of the region pairs showed statistically significant sex differences in covariance (**Figure 2 B-D**: medial amygdala – BNST: Δ = 0.053, p = 1.0; olfactory bulb – BNST: Δ = 0.041, p = 1.0; medial amygdala – olfactory bulb: Δ = 0.083, p = 1.0).

**Figure 2.**
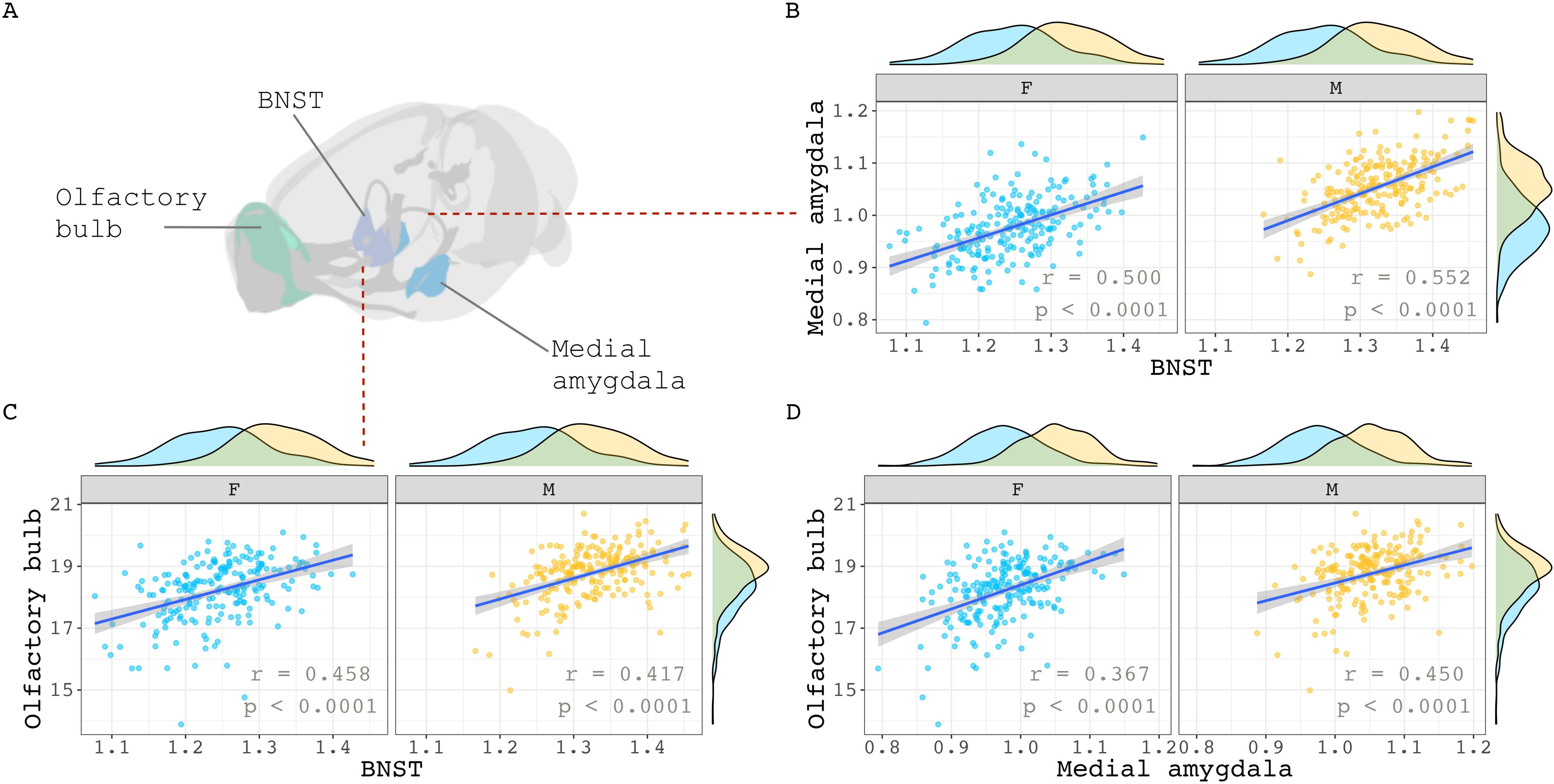
Within sex correlations between classic sex-biased regions in the mouse. **(A)** Visualization of three volumetrically sex-biased regions: olfactory bulb, BNST, and medial amygdala. **B, C, D)** Sex-specific correlations of the medial amygdala with BNST (B), olfactory bulb with BNST (C), and olfactory bulb with medial amygdala (D). Sex-specific means of each region were added to their residuals before correlation calculations began. This was done to maintain volumetric sex differences in volume distribution for visualization purposes. Correlation sex differences for all three pairs were not statistically significant (p = 1.0).

Second, we expanded our analyses to characterize the relationship between sex-biased volume and sex-biased volume covariance throughout the brain of each species more broadly. To contextualize these analyses, we projected all observed sex-differences in volume covariance into anatomical space by computing the mean signed sex-difference in volume covariance for each region and visualizing the distribution of this regional value across the brain of each species (**Figure 3 A, B** – left columns). Re-computing these maps using information for just those region pairs with statistically significant sex differences in covariance (see Methods) highlighted several brain regions in each species with significant cumulative sex-differences in volume covariance with the rest of the brain (**Figure 3 A, B** -- right columns). In mice, these regions had almost exclusively stronger covariance in females and included the infralimbic area, medial parietal association cortex, and the pons. Regional covariance sex differences were also more often female-biased in humans and included regions such as the auditory complex 4, the primary sensory cortex, and the primary motor cortex. However, in humans, we also observed regions with significantly male-biased sex-differences in regional structural covariance, including the area prostriata and the premotor eye field. Qualitatively some of the regions highlighted by these analyses also showed sex-differences in mean volume [e.g. mice: mamillary body, mouth primary somatosensory area, claustrum (female-biased mean volume); BNST, olfactory bulb, CA3 pyramidal region (male-biased volume) / humans: primary sensory cortex, cingulate regions (supplementary and cingulate eye field, ventral area 24d) (female-biased volume); hypothalamus, amygdala, posterior insular region 1 (male-biased volume). The full lists of mean regional covariance sex differences in each species can be found in **Extended Tables 3-1, 3-2**.

**Figure 3.**
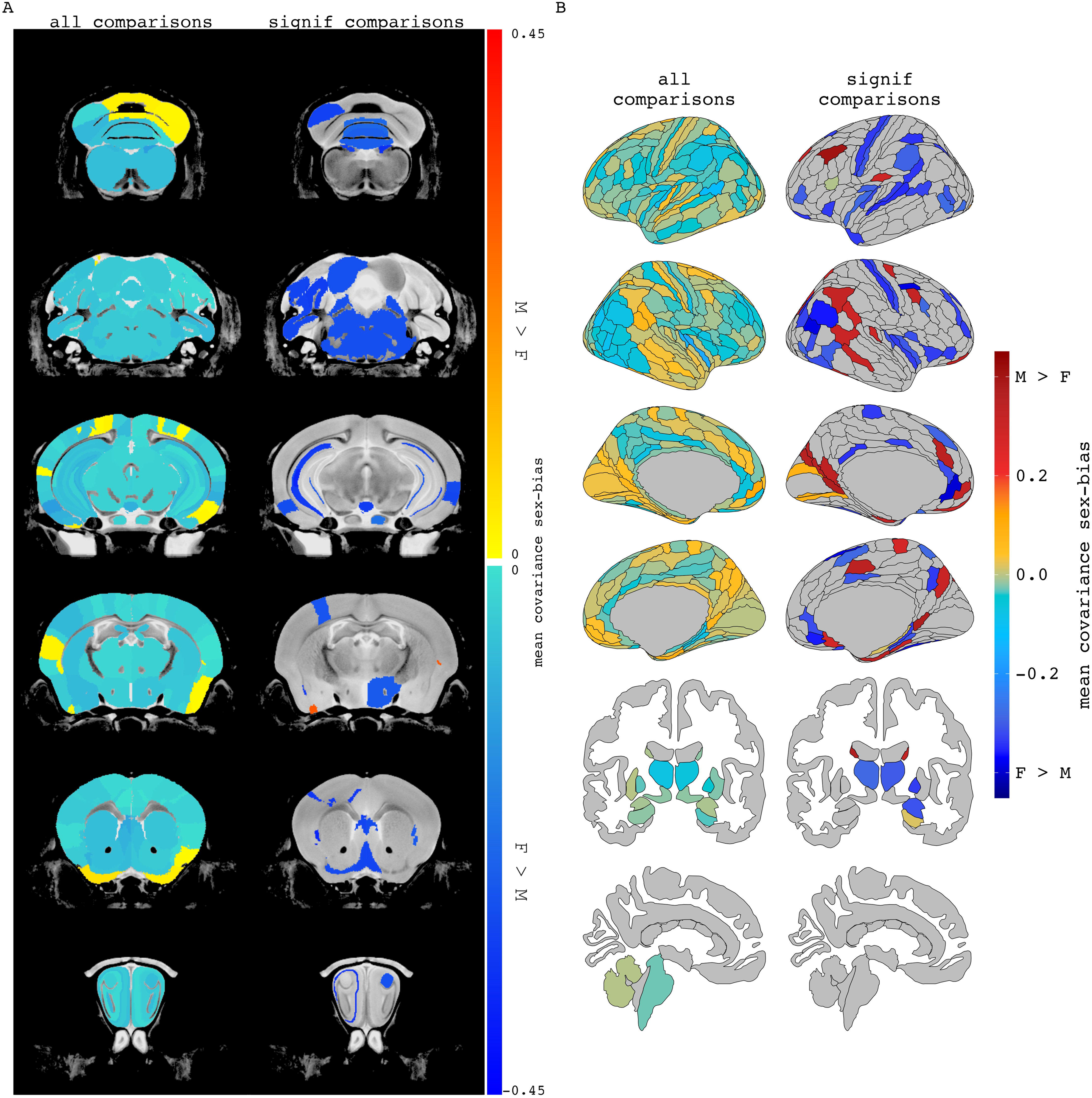
Regional mean covariance sex differences in mice and humans. Mean covariance sex differences in mouse (A) and human (B) when averaging a region’s covariance sex difference across all pairwise comparisons (all comparisons) or only pairwise comparisons with statistically significant covariance sex differences (significant comparisons). When including all pairwise comparisons, 89% of mouse regions had mean covariance sex differences which were stronger in females. For humans, averaging all pairwise covariance sex differences per brain region resulted in 68% of human regions with female-biased mean covariance. When including only comparisons significant for covariance sex differences, 96% of the included mouse regions and 70% of the human regions were female biased. The magnitude and direction of sex-bias of left and right regions generally correlate when considering all sex differences results (mice: r = 0.57; humans: r = 0.48).

To disentangle individual inter-regional pairs from these regional summaries of Figure 3, and to further assess the involvement of volumetrically sex-biased regions in sex-biased covariance patterns, we generated graphs containing all inter-regional pairs with statistically significant sex- biased volume covariance in each species (regions as nodes and sex-biased covariance relationships as edges). **Figures 4 and 5** represent the largest connected components of these graphs for mice and humans. In mice, the largest components are centered around the right cuneate nucleus and the left infralimbic area. All connecting covariance sex-biased edges are female-biased. Of the 27 regions involved in these components, 7 volumetrically sex-biased regions are distantly associated with the central nodes (nodes with 3 or more associated edges through intermediary connections). These regions include the male-biased hypothalamus and CA3 pyramidal regions, and the female-biased mamillary body and the cerebellar crus regions. The largest connected component in humans contains 35 regions and is mainly centered around the right posterior insular area 1, a volumetrically male-biased region, and the right parainsular region area 52. Approximately 80% of this component’s edges are female-biased. Only 2 other volumetrically sex-biased regions are represented in this component (supplementary and cingulate eye field; primary sensory cortex – both are female-biased).

**Figure 4.**
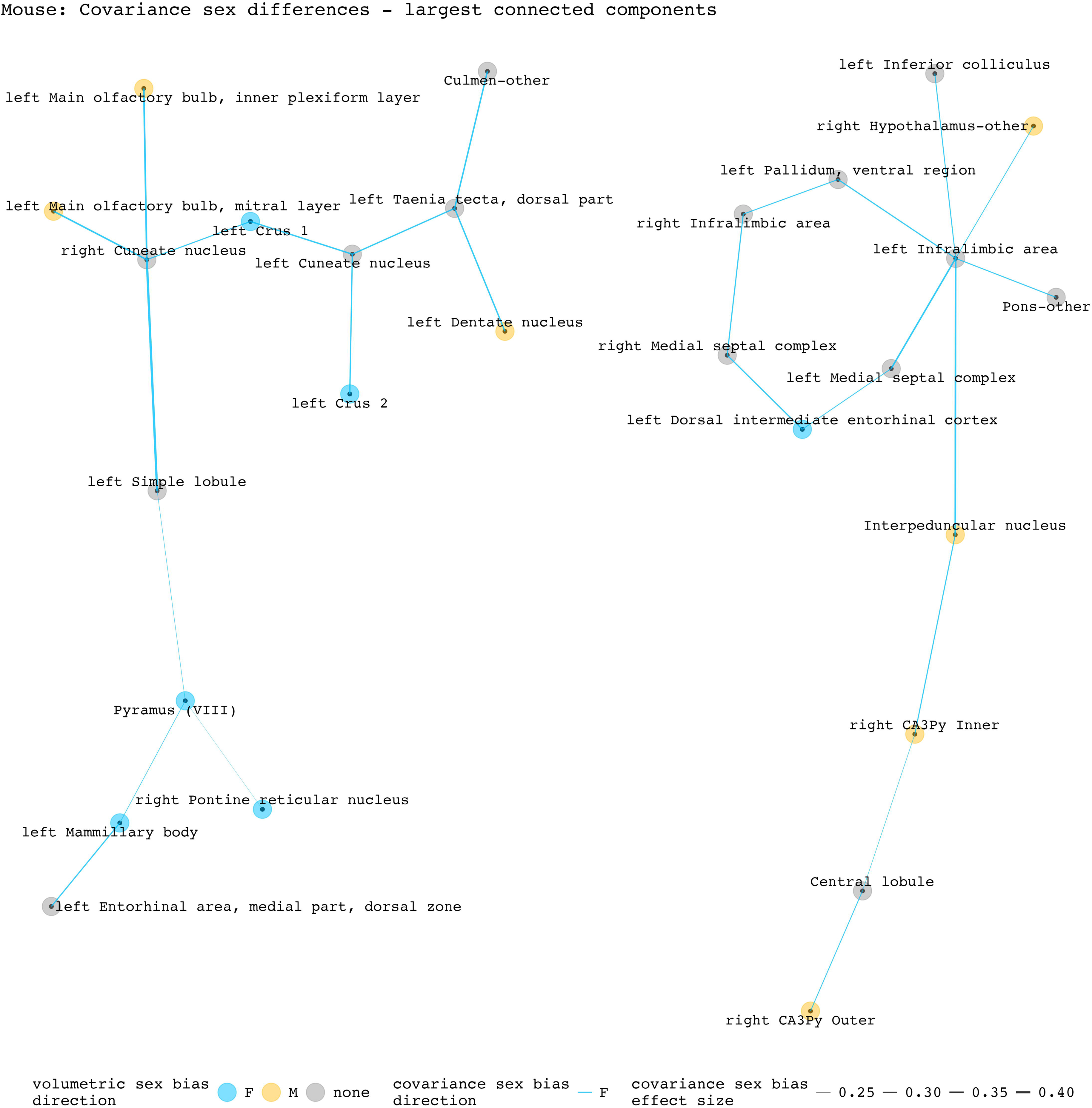
Statistically significant covariance sex differences in mice, represented as nodes and edges (top 2 most connected components). The first component (left side) is centered by the right cuneate nucleus and involves 14 regions/13 covariance pairs. The second component (right side) is centered on the left infralimbic area and involves 13 regions/13 covariance pairs. Both components contain only female-biased edges. Of the 16 edges involving volumetrically sex-biased regions in these components, 13 are between a volumetrically sex-biased region and a region without any volumetric sex bias; 3 are between two volumetrically sex-biased regions.

**Figure 5.**
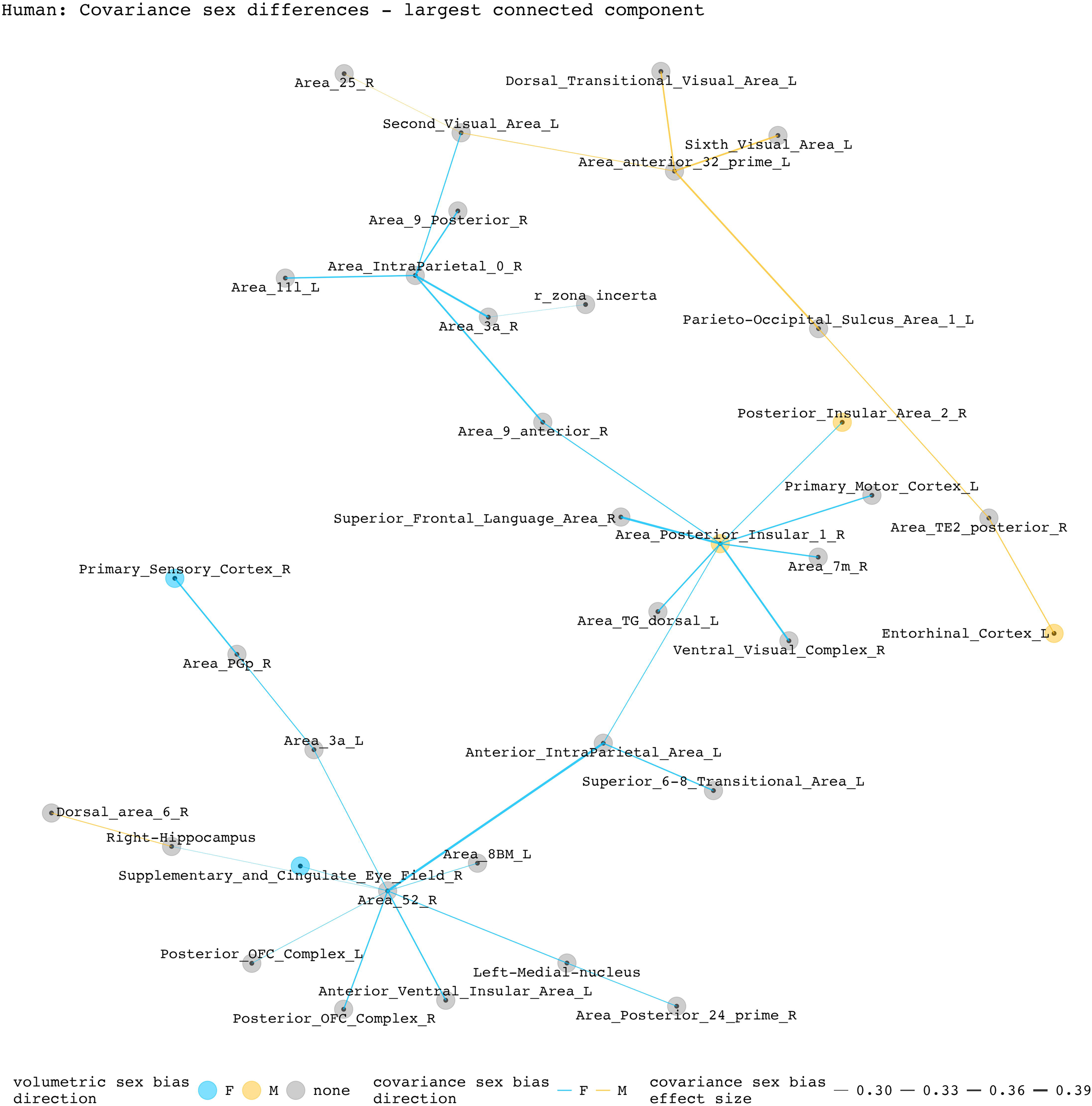
Statistically significant covariance sex differences in humans, represented as nodes and edges (most connected component). This component is centered by two regions: right parainsular area 52 with 7 connecting nodes and right posterior insular area 1 with 8 connecting nodes. Of the 33 edges included in this component, 8 are male-biased. Of the 35 regions included in this component, 3 have volumetric sex differences (right posterior insular area 1, right posterior insular area 2, right supplementary and cingulate eye field). Except for the female-biased covariance relationship between the right posterior insular area 1 and the posterior insular area 2, all sex-biased edges involving a volumetrically sex-biased region (9 in total) are paired with a volumetrically non-sex-biased region.

Finally, we sought to quantitatively test – within both species - if regional variation in the magnitude of sex-biased volume covariance was related to regional variation in the magnitude of sex-biased mean volume. These analyses were repeated twice within each species - once using absolute values and once using signed values for mean regional sex differences in volume covariance (absolute inter-regional sex differences in covariance for computing regional means and absolute sex differences in volume) for all available regions. The absolute values analysis shows no evidence of an association between regional volumetric and mean covariance sex-bias effect size in both humans and mice (**Fig. 6A, B**, mice: r = 0.03, p = 0.68; humans: r = 0.13, p = 0.98). However, the signed analyses revealed that both species show a weak yet statistically significant inverse relationship between regional covariance and volumetric sex-bias directions (**Fig. 6 C, D,** mice: r = -0.19, p = 0.002; humans: r = -0.19; p = 0.001). Specifically, in both species, regions of significantly male-biased volume more often show female-biased volume covariance and vice versa for regions of significantly female-biased volume.

**Figure 6.**
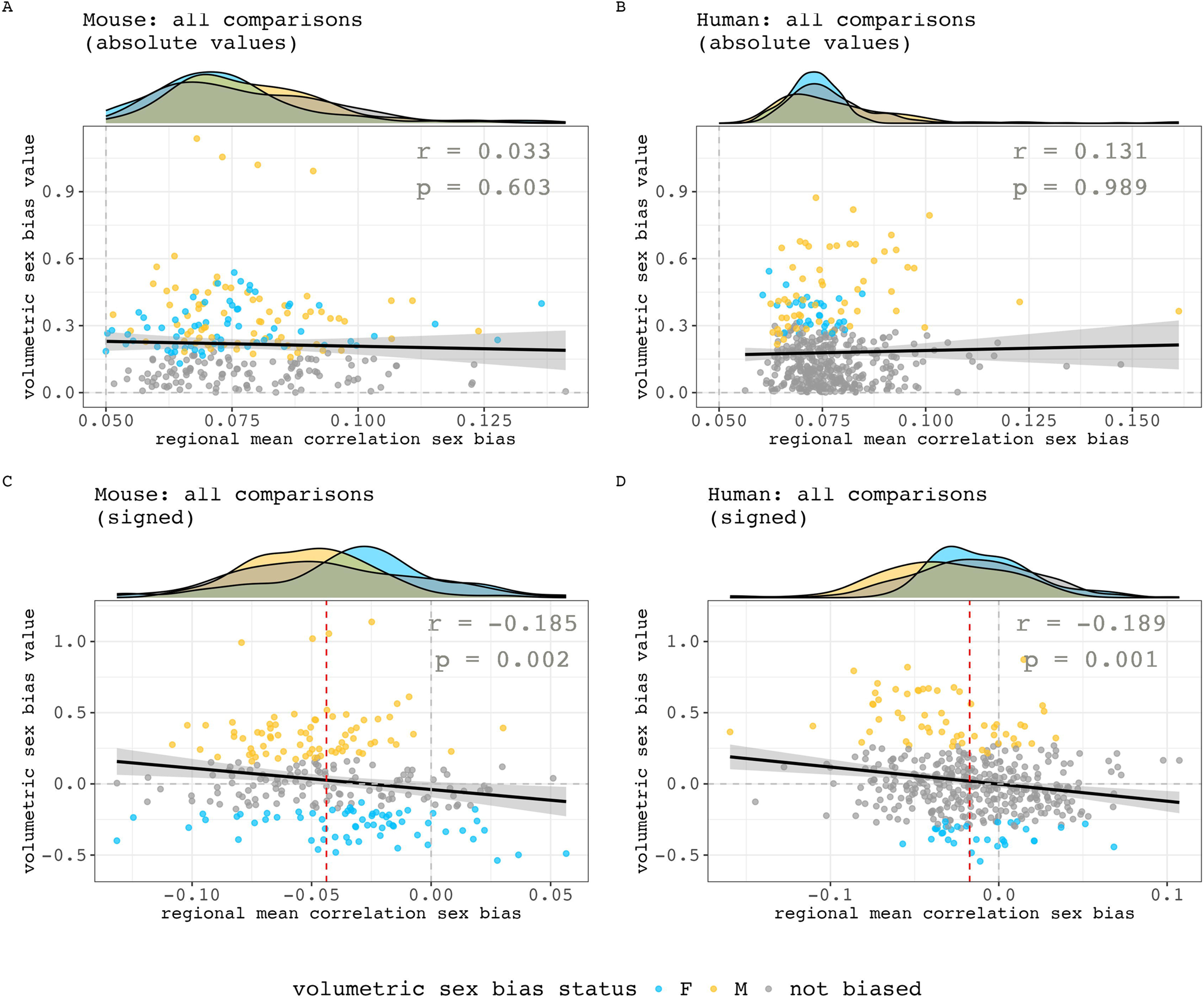
Relationships between the sex-biased in regional volumetric and regional volume covariance. **A, B)** Relationships between absolute values in mean regional correlation sex bias and volumetric sex biases in mouse (A) and human (B). **C, D**) Relationships between signed values in mean regional correlation sex bias and volumetric sex biases in mouse (C) and human (D). Negative values are female biased in both axes. Marginal density plots represent the regional mean correlation sex-bias distributions of different volumetric sex-bias categories (female-biased, male-biased and not significantly sex-biased). For the signed analyses in both species, there is a statistically significant, but weak negative correlation between sex-differences in volume and sex-differences in volume covariance (inset statistics). Thus - in both species - regions of significantly male-biased volume more often show female-biased volume covariance and vice versa for regions of significantly female-biased volume. However, note that regions with the largest volumetric sex-bias values do not have the largest mean correlation sex-bias – their values are concentrated closer to the median regional mean correlation sex bias (red dashed line). Also, regions with statistically significant volumetric sex biases but smaller volumetric sex bias values can be found on either end of the x-axis.

Taken together, these analyses help to localize sex-differences in volume covariance within the brains of humans and mice and further specify the spatial relationship between this phenomenon and accompanying sex-differences in regional volume. We find several regions of prominently sex-biased volume covariance in each species – highlighting the infralimbic area, medial parietal cortex, and pons in mice and auditory complex 4, the primary sensory cortex, and primary motor cortex in humans. Some of these regions overlap with regional sex-differences in volume – highlighting brain areas that show two forms of sex-biased organization (e.g. mamillary body, mouth primary somatosensory area, claustrum in mouse, hypothalamus, amygdala, and posterior insular region 1 in humans). Although significant sex differences in anatomical covariance are not seen amongst major classical foci of sex-biased volume in the mouse brain, there is a subtle yet significant inverse association between sex differences in volume and volume covariance across both the murine and human brain.

## DISCUSSION

In this study, we utilize sex as a natural experiment to ask whether developmental programming affects structural covariance formation within humans and mice. Our results provide a systematic survey of sex-biased neuroanatomical covariance and their conservation across species. We consider each of our main findings below.

First, en route to estimating sex-differences in structural covariance across species, we observe that mice show stronger neuroanatomical covariance than humans. This observation might reflect the combined action of several species’ differences. Greater genetic and environmental variability across humans compared to inbred laboratory mice likely translates into weaker coordination of anatomical variation within human brain. Greater neuroanatomical covariation in mice may also track with the gross brain size differences between species – larger vertebrates tend to show greater phenotypic variability (Hallgrímsson and Maiorana, 2000) and inter- regional covariance strength likely drops off with the greater inter-regional distances within the human compared to the murine brain (He et al., 2007; Yee et al., 2018). Species differences in overall neuroanatomical covariance strength could coincide with substantially longer lifespan in humans than mice given evidence of age-related decreases in neuroanatomical covariance within humans (DuPre and Spreng, 2017).

Second, we replicate prior reports of female-biased volume covariance in the human brain (Wierenga et al., 2022; Shi et al., 2023) and reveal that this is also evident in the mouse brain. This species convergence suggests that female-biased neuroanatomical covariance may be an evolutionarily conserved feature, though studies in other species is required to verify this. Many previously proposed candidate mechanisms for sex-biased anatomical covariance cannot parsimoniously account for the female-biased covariance observed in both species. Although sex-differences in overall brain-size would predict stronger anatomical covariation in female humans (given larger brain size in males) – this explanation could not account for sex-biased anatomical covariation in mice as they do not have overall brain size sex differences (Guma et al., 2024). Similarly, although greater anatomical variability in human males could explain stronger anatomical covariation in human females (if male-biased anatomical variance in humans reflects greater developmental noise that is uncorrelated between regions), this explanation would not apply in mice – which do not show the prominent sex-bias in neuroanatomical variability evident in humans (Wierenga et al., 2018; Wierenga et al., 2022; Guma et al., 2024). A larger human study also suggests that biological variation is a poor predictor of covariation strength: although subcortical volumes and cortical thickness are more variable in males, only cortical thickness correlations were higher in males (Wierenga et al., 2022). The most parsimonious hypothesis for our findings would be that the tendency towards female-biased neuroanatomical covariance in humans and mice reflects a genetic and hormonal aspects of sex shared between species and capable of shaping inter-regional anatomical covariance. For example, females are biallelic for X-linked gametologs and males are hemizygous for X- and Y-member of each gametolog pair in both species. Given this, regionally specific functional divergences of X- vs. Y-gametologs (DeCasien et al., 2024) could represent a male-specific source of inter-regional divergence in neurobiological organization operative in both species, leading to stronger inter-regional covariance in females.

Third, we profile the regional distribution of sex-biased volume covariance in each species and probe how it relates to sex differences in mean regional volume. As an initial targeted test for the idea of coordinated sex-differences in brain volume and covariance – we focused on 3 canonically sex-biased regions in the mouse brain (BNST, medial amygdala, olfactory bulb) and did not find sex-biased volumetric covariance amongst these regions. Thus, male-specific processes driving highly reproducible male-bias in these regions’ mean volumes (Williams et al., 2001; Forger et al., 2004; Morris et al., 2008) do not appear to open male-specific sources of covariation between regions. However, we cannot rule out that these regions have sex-specific volume covariation sources that counterbalance each other or are hidden by dominantly shared sources of inter-regional volume covariation between the sexes.

Fourth, while no covariance sex differences were identified among classical regions of volumetric sex bias, broadening the search for sex-biased volume covariance identified multiple brain regions showing differential brain wide volume coupling between sexes. Echoing the female-bias volume covariance found in our global analyses, regional sex-biased covariance hotspots were almost exclusively female-biased in mice and mostly female-biased in humans. Regions with significant sex-bias in covariance in mice were the infralimbic area, medial parietal association cortex, and the pons (stronger in females); in humans, these regions include the auditory complex 4, the primary sensory cortex, the primary motor cortex (stronger in females), the area prostriata, and the premotor eye field (stronger in males). Some of the brain regions showing prominent sex-biased covariance are also volumetrically sex-biased in our study, such as the mamillary body and claustrum in mice and the posterior insular region 1 and hypothalamus in humans. These regions offer high-priority targets for follow-up mechanistic analyses and for probing how sex-biased anatomical organization of the brain might relate to brain function.

Fifth, to identify specific covariance sex differences that underpin these regional patterns, we visualized networks of significant covariance sex differences in mice and humans, flagging large sets of brain regions interlinked through sex-biased volume covariance. Some of the pairings in these networks have been shown through tract tracing and diffusion tensor MRI studies to be physically connected (AIBS:#177889243; Oh et al., 2014; Ghaziri et al., 2015; Harris et al., 2019). These findings help motivate challenging experimental studies that will now be needed to determine the drivers for sex-differences in neuroanatomical covariance, such as sex differences in structural connectivity.

Finally, while we define some instances of overlap between sex-biased volume covariance and sex-biased volume, we show that these features are not strongly related across the brain. This dissociation implies that processes regulating sex-biased volume and sex-biased covariance development are largely dissociable. However, we do observe a weak negative correlation between signed sex differences in brain volume and brain volume covariance – mostly driven by a tendency for volumetrically male-biased regions to show female-biased volume covariance. Thus, we speculate that male-specific influences on the size of a sex-biased region introduces male-specific sources of regional volume variation that are poorly coordinated between regions – resulting in weakened correlations amongst these regions in males.

Our findings should be considered with several limitations. First – we focused on brain volume as a phenotype that can be compared across species by sMRI. However, sMRI can only resolve sex differences in volume and covariance at or above the lower limit of its voxel dimensions (here, ∼ 40 µm^3^ in mice and ∼1 mm^3^ in humans). Sex differences in structures indiscernible by contrast in T2-weighted MRI are also not detectable. Second, there are brain properties beyond volume, such as regional myelin content and cortical thickness. We therefore cannot assume our volumetric findings will generalize to other properties. Third, we estimate covariance cross-sectionally in adulthood datasets in each species. As such, we cannot speak to age-related variations of sex-biased anatomical covariance. Fourth, we note that there is not straightforward convergence between brain regions that show prominent sex-differences in volume covariance in humans versus mice (Figures 4-5). However, theoretical and methodological approaches for inferring homologies between human and mouse brains are rapidly evolving (Beauchamp et al., 2022; Guma et al., 2024) and could be applied in the future to formally assess regional species convergence. Finally, our study design is purely observational, and we cannot dissect potential mechanistic or functional bases for observed sex differences in covariance. This is especially true given the maximum magnitude in correlation differences is small and there are few covariance sex differences overall. In mice, these differences could ultimately be related to sex-biased chromosomal and gonadal influences on brain development. Such influences are likely operative in humans but will certainly be interwoven with co-occurring gendered influences stemming from the external environment. Transgenic models like the Four Core Genotypes could explain how covariance sex differences are formed (Arnold and Chen, 2009; Corre et al., 2016). These experiments would expand our understanding of how sex affects brain organization and how developmental programming can drive covariance formation.

Notwithstanding the limitations above, our study design was able to use sex differences as an informative probe for theories regarding the developmental bases of neuroanatomical covariance, and – en route to doing so - provide a thorough parallel mapping of global and inter-regional sex differences in brain volume covariance for the human and murine brain. We validate existing human results that show females have stronger covariance and show the same phenotype in mice. We provide a fine-grained delineation of brain systems that show sex-biased covariance in each species. Our finding that structural covariance sex differences minimally involve volumetrically sex-biased structures supports the viewpoint that the sex influences likely have uncoordinated effects across brain regions and highlights structural covariance as a novel axis of sex-biased brain organization warranting further study. Additionally, the cross-species approach used here allows us to articulate the potential impacts of stochasticity and variation on structural covariance, emphasizing the importance of synchronizing methodologies across species for gaining new insights into brain biology and increasing the translatability of animal findings (Barron et al., 2021).

## Acknowledgements

This research was supported by the Intramural Research Program of the NIMH (NIH annual report number ZIAMH002949-08), the National Institute of Child Health and Disease (RO1HD100298), The Canadian Institutes of Health Research, BrainCanada, the Ontario Brain Institute. The Wellcome Centre for Integrative Neuroimaging is supported by core funding from the Wellcome Trust (203139//16/Z and 203139/A/16/Z). Computational support for this research was provided by Compute Canada (https://computecanda.ca) and the NIH HPC Biowulf cluster (https://hpc.nih.gov). LP is an MD/PhD student supported by the NIH Oxford-Cambridge Scholars Program and the South Texas MSTP (GM145432-02). For the purpose of open access, the author has applied a CC BY public copyright license to any Author Accepted Manuscript version arising from this submission.

## EXTENDED FIGURES

**Figure 1-1. Male and female structural covariance of mouse.** Pearson correlation matrices of regional brain volumes in mice, separated by sex. Each row and column represent a grey matter structure defined by the Allen Mouse Brain Atlas, shown in the right column. Structures are denoted by color bands that correspond to their structures’ colors in the atlas. Each element in the matrices represent a Pearson correlation between two brain structures. Mouse brain structures mainly have positive correlations to each other. Given the same correlation color scale across sex, female mice appear to have more intense positive correlations in the brain comparing to males.

**Figure 1-2. Male and female structural covariance of human.** Pearson correlation matrices of regional brain volumes in humans, separated by sex. Cortical structures are grouped into 6 lobes, as defined by Freesurfer output for Glasser atlas segmentation: occipital (light blue), frontal (dark blue), parietal (light green), temporal (dark green), insula (salmon). Subcortical structures are defined as followed using Freesurfer output for aseg atlas segmentation: thalamus proper (red), amygdala (orange), basal ganglia – combination of caudate, putamen, pallidum (light purple), pons (dark purple), cerebellum (yellow), ventral diencephalon (brown). Additional segmentations of amygdala and hypothalamic nuclei were grouped under the amygdala and ventral diencephalon categories, respectively. Structures are denoted by color bands that correspond to their structures’ colors in the atlas. Each element in the matrices represenst a Pearson correlation between two brain structures. Human brain structures mainly have positive correlations to each other. Given the same correlation color scale across sex, females appear to have more intense positive correlations in the brain comparing to males. The correlation color intensities are less than those observed in both male and female mice

**Figure 1-3. Simulated data demonstration: Shared versus sex-specific developmental influences on structural covariance sex differences**. The correlation strength between two structures tends to increase as a function of shared developmental influences, such as through shared axonal connectivity or gene expressions. As structures share less influences, their correlations also tend to weaken. If sex-specific influences only act on certain structures in the brain, then they are more likely to influence the covariance between pairs where one structure receives the developmental influences of sex while the other does not. In other words, sex-specific developmental influences are more likely to act upon structure pairs with less shared influences to each other, or pairs with weaker correlations. For this reason, one could expect covariance sex differences to be largest between covariance pairs with weak associations to each other

**Figure 1-4. Structural covariance strength versus sex difference in mice and humans. A-D)** Pairwise Pearson correlations versus absolute correlation sex differences for all comparisons in female mouse (A) and human (B) and male mouse (C) and human (D). **E, F)** Pairwise Pearson correlations versus absolute correlation sex differences at different covariance sex differences significance thresholds for mouse (E) and human (F). Association strengths between the covariance sex differences of significant pairs and their Pearson correlations are calculated using Pearson correlation and p-values generated by the cor.test function in R. The predicted inverse relationship between covariance strength and sex differences are more prominent in mice than humans.

## Extended Tables Captions

**Extended Table 1-1.** All pairwise covariance sex difference results in mice. Differences significance status is defined in the “signif” and “signif_adj” columns, corresponding to whether the difference is significant after permutation testing or after permutation testing and multiple comparisons corrections.

**Extended Table 1-2.** All pairwise covariance sex difference results in humans. Differences significance status is defined in the “signif” and “signif_adj” columns, corresponding to whether the difference is significant after permutation testing or after permutation testing and multiple comparisons corrections.

**Extended Table 3-1.** All regional mean covariance sex differences in mice. Signed values are indicated as “meanCov_signed”. Absolute values are indicated as “meanCov_absolute”.

**Extended Table 3-2.** All regional mean covariance sex differences in humans. Signed values are indicated as “meanCov_signed”. Absolute values are indicated as “meanCov_absolute”.

## REFERENCES

AIBS:#177889243 Allen Institute for Brain Science. Experiment 177889243. In. Allen Institute for Brain Science.

Alexander-Bloch A, Raznahan A, Bullmore E, Giedd J (2013) The Convergence of Maturational Change and Structural Covariance in Human Cortical Networks. Journal of Neuroscience 33:2889.

Andrews TJ, Halpern SD, Purves D (1997) Correlated Size Variations in Human Visual Cortex, Lateral Geniculate Nucleus, and Optic Tract. Journal of Neuroscience 17:2859–2868.

Arnold AP, Chen X (2009) What does the “four core genotypes” mouse model tell us about sex differences in the brain and other tissues? Front Neuroendocrinol 30:1–9.

Avants B, Tustison NJ, Song G (2009) Advanced Normalization Tools: V1.0. Insight Journal.

Avants BB, Tustison NJ, Song G, Cook PA, Klein A, Gee JC (2011) A reproducible evaluation of ANTs similarity metric performance in brain image registration. NeuroImage 54:2033–2044.

Barron HC, Mars RB, Dupret D, Lerch JP, Sampaio-Baptista C (2021) Cross-species neuroscience: closing the explanatory gap. Philosophical Transactions of the Royal Society B 376:20190633.

Beauchamp A, Yee Y, Darwin BC, Raznahan A, Mars RB, Lerch JP (2022) Whole-brain comparison of rodent and human brains using spatial transcriptomics. eLife 11:e79418.

Benjamini Y, Hochberg Y (1995) Controlling the False Discovery Rate: A Practical and Powerful Approach to Multiple Testing. Journal of the Royal Statistical Society: Series B (Methodological) 57:289–300.

Benjamini Y, Yekutieli D (2001) The control of the false discovery rate in multiple testing under dependency. Annals of Statistics 29.

Cahill LS, Laliberté CL, Ellegood J, Spring S, Gleave JA, van Eede MC, Lerch JP, Henkelman RM (2012) Preparation of fixed mouse brains for MRI. NeuroImage 60:933–939.

Chakravarty MM, Steadman P, van Eede MC, Calcott RD, Gu V, Shaw P, Raznahan A, Collins DL, Lerch JP (2013) Performing label-fusion-based segmentation using multiple automatically generated templates. Human Brain Mapping 34:2635–2654.

Chung MK, Worsley KJ, Paus T, Cherif C, Collins DL, Giedd JN, Rapoport JL, Evans AC (2001) A Unified Statistical Approach to Deformation-Based Morphometry. NeuroImage 14:595–606.

cocoframer (2018) © 2018 Allen Institute for Brain Science. Allen Brain Explorer. Available from: connectivity.brain-map.org/3d-viewer/. In. Allen Institute for Brain Science.

Collins DL, Neelin P, Peters TM, Evans AC (1994) Automatic 3D intersubject registration of MR volumetric data in standardized Talairach space. Journal of Computer Assisted Tomography 18:192–205.

Corre C, Friedel M, Vousden DA, Metcalf A, Spring S, Qiu LR, Lerch JP, Palmert MR (2016) Separate effects of sex hormones and sex chromosomes on brain structure and function revealed by high-resolution magnetic resonance imaging and spatial navigation assessment of the Four Core Genotype mouse model. Brain Struct Funct 221:997–1016.

Dale A, Fischl B, Sereno MI (1999) Cortical Surface-Based Analysis: I. Segmentation and Surface Reconstruction. NeuroImage 9:179–194.

Dale AM, Sereno MI (1993) Improved Localizadon of Cortical Activity by Combining EEG and MEG with MRI Cortical Surface Reconstruction: A Linear Approach. Journal of Cognitive Neuroscience 5:162–176.

DeCasien AR, Guma E, Liu S, Raznahan A (2022) Sex differences in the human brain: a roadmap for more careful analysis and interpretation of a biological reality. Biology of Sex Differences 13:43.

DeCasien AR, Tsai K, Liu S, Thomas A, Raznahan A (2024) Evolutionary divergence between homologous X-Y chromosome genes shapes sex-biased biology. Nature Ecology and Evolution.

Desikan RS, Ségonne F, Fischl B, Quinn BT, Dickerson BC, Blacker D, Buckner RL, Dale AM, Maguire RP, Hyman BT, Albert MS, Killiany RJ (2006) An automated labeling system for subdividing the human cerebral cortex on MRI scans into gyral based regions of interest. NeuroImage 31:968–980.

Devenyi G (2024) Library of Bpipe functions for processing Minc files, version c7561d6. In.

Dorr AE, Lerch JP, Spring S, Kabani N, Henkelman RM (2008) High resolution three-dimensional brain atlas using an average magnetic resonance image of 40 adult C57Bl/6J mice. NeuroImage 42:60–69.

DuPre E, Spreng RN (2017) Structural covariance networks across the life span, from 6 to 94 years of age. Netw Neurosci 1:302–323.

Ellegood J et al. (2015) Clustering autism: using neuroanatomical differences in 26 mouse models to gain insight into the heterogeneity. Molecular Psychiatry 20:118–125.

Eskildsen SF, Coupé P, Fonov V, Manjón JV, Leung KK, Guizard N, Wassef SN, Østergaard LR, Collins DL (2012) BEaST: Brain extraction based on nonlocal segmentation technique. NeuroImage 59:2362–2373.

Fischl B (2012) FreeSurfer. NeuroImage 62:774–781.

Fischl B, Dale AM (2000) Measuring the thickness of the human cerebral cortex from magnetic resonance images. Proc Natl Acad Sci U S A 97:11050–11055.

Fischl B, Sereno MI, Dale A (1999a) Cortical Surface-Based Analysis: II: Inflation, Flattening, and a Surface-Based Coordinate System. NeuroImage 9:195 - 207.

Fischl B, Liu A, Dale AM (2001) Automated manifold surgery: constructing geometrically accurate and topologically correct models of the human cerebral cortex. IEEE Transactions on Medical Imaging 20:70–80.

Fischl B, Sereno MI, Tootell RBH, Dale AM (1999b) High-resolution intersubject averaging and a coordinate system for the cortical surface. Human Brain Mapping 8:272–284.

Fischl B, Dale AM, Sereno MI, Tootell RBH, Rosen BR (1998) A Coordinate System for the Cortical Surface. Neuroimage 7:S740.

Fischl B, Salat DH, van der Kouwe AJW, Makris N, Ségonne F, Quinn BT, Dale AM (2004a) Sequence-independent segmentation of magnetic resonance images. NeuroImage 23:S69–S84.

Fischl B, Salat DH, Busa E, Albert M, Dieterich M, Haselgrove C, van der Kouwe A, Killiany R, Kennedy D, Klaveness S, Montillo A, Makris N, Rosen B, Dale AM (2002) Whole brain segmentation: automated labeling of neuroanatomical structures in the human brain. Neuron 33:341–355.

Fischl B, van der Kouwe A, Destrieux C, Halgren E, Ségonne F, Salat DH, Busa E, Seidman LJ, Goldstein J, Kennedy D, Caviness V, Makris N, Rosen B, Dale AM (2004b) Automatically Parcellating the Human Cerebral Cortex. Cereb Cortex 14:11–22.

Forger NG, Rosen GJ, Waters EM, Jacob D, Simerly RB, de Vries GJ (2004) Deletion of Bax eliminates sex differences in the mouse forebrain. Proc Natl Acad Sci U S A 101:13666–13671.

Fortin J-P, Parker D, Tunç B, Watanabe T, Elliott MA, Ruparel K, Roalf DR, Satterthwaite TD, Gur RC, Gur RE, Schultz RT, Verma R, Shinohara RT (2017) Harmonization of multisite diffusion tensor imaging data. Neuroimage 161:149–170.

Fortin J-P, Cullen N, Sheline YI, Taylor WD, Aselcioglu I, Cook PA, Adams P, Cooper C, Fava M, McGrath PJ, McInnis M, Phillips ML, Trivedi MH, Weissman MM, Shinohara RT (2018) Harmonization of cortical thickness measurements across scanners and sites. NeuroImage 167:104–120.

Friedel M, van Eede MC, Pipitone J, Chakravarty MM, Lerch JP (2014) Pydpiper: a flexible toolkit for constructing novel registration pipelines. Frontiers in Neuroinformatics 8.

Galton F (1888) Co-relations and Their Measurement. Proceedings of the Royal Society of London:135–145.

Ge R, Liu X, Long D, Frangou S, Vila-Rodriguez F (2021) Sex effects on cortical morphological networks in healthy young adults. NeuroImage 233:117945.

Ghaziri J, Tucholka A, Girard G, Houde J-C, Boucher O, Gilbert G, Descoteaux M, Lippé S, Rainville P, Nguyen DK (2015) The Corticocortical Structural Connectivity of the Human Insula. Cerebral Cortex 27:1216–1228.

Glasser MF, Sotiropoulos SN, Wilson JA, Coalson TS, Fischl B, Andersson JL, Xu J, Jbabdi S, Webster M, Polimeni JR, Van Essen DC, Jenkinson M (2013) The minimal preprocessing pipelines for the Human Connectome Project. Neuroimage 80:105–124.

Glasser MF, Coalson TS, Robinson EC, Hacker CD, Harwell J, Yacoub E, Ugurbil K, Andersson J, Beckmann CF, Jenkinson M, Smith SM, Van Essen DC (2016) A multi-modal parcellation of human cerebral cortex. Nature 536:171–178.

Glur C (2020) data.tree: General Purpose Hierarchical Data Structure. In.

Gong G, He Y, Chen ZJ, Evans AC (2012) Convergence and divergence of thickness correlations with diffusion connections across the human cerebral cortex. NeuroImage 59:1239–1248.

Gorski RA, Gordon JH, Shryne JE, Southam AM (1978) Evidence for a morphological sex difference within the medial preoptic area of the rat brain. Brain Research 148:333–346.

Guma E, Beauchamp A, Liu S, Levitis E, Ellegood J, Pham L, Mars RB, Raznahan A, Lerch JP (2024) Comparative neuroimaging of sex differences in human and mouse brain anatomy. eLife 13.

Hallgrímsson B, Maiorana V (2000) Variability and size in mammals and birds. Biological Journal of the Linnean Society 70:571–595.

Han X, Jovicich J, Salat D, van der Kouwe A, Quinn B, Czanner S, Busa E, Pacheco J, Albert M, Killiany R, Maguire P, Rosas D, Makris N, Dale A, Dickerson B, Fischl B (2006) Reliability of MRI-derived measurements of human cerebral cortical thickness: The effects of field strength, scanner upgrade and manufacturer. NeuroImage 32:180–194.

Harris JA et al. (2019) Hierarchical organization of cortical and thalamic connectivity. Nature 575:195–202.

He Y, Chen ZJ, Evans AC (2007) Small-World Anatomical Networks in the Human Brain Revealed by Cortical Thickness from MRI. Cereb Cortex 17:2407–2419.

Hines M, Allen LS, Gorski RA (1992) Sex differences in subregions of the medial nucleus of the amygdala and the bed nucleus of the stria terminalis of the rat. Brain Research 579:321–326.

Iglesias JE, Van Leemput K, Bhatt P, Casillas C, Dutt S, Schuff N, Truran-Sacrey D, Boxer A, Fischl B (2015a) Bayesian segmentation of brainstem structures in MRI. NeuroImage 113:184–195.

Iglesias JE, Augustinack JC, Nguyen K, Player CM, Player A, Wright M, Roy N, Frosch MP, McKee AC, Wald LL, Fischl B, Van Leemput K (2015b) A computational atlas of the hippocampal formation using ex vivo, ultra-high resolution MRI: Application to adaptive segmentation of in vivo MRI. NeuroImage 115:117–137.

Johnson WE, Li C, Rabinovic A (2007) Adjusting batch effects in microarray expression data using empirical Bayes methods. Biostatistics 8:118–127.

Jovicich J, Czanner S, Greve D, Haley E, van der Kouwe A, Gollub R, Kennedy D, Schmitt F, Brown G, MacFall J, Fischl B, Dale A (2006) Reliability in multi-site structural MRI studies: Effects of gradient non-linearity correction on phantom and human data. NeuroImage 30:436–443.

Kuhn M, Wickham H (2020) Tidymodels: a collection of packages for modeling and machine learning using tidyverse principles. In.

Kuperberg GR, Broome M, McGuire PK, David AS, Eddy M, Ozawa F, Goff D, West WC, Williams SCR, van der Kouwe A, Salat D, Dale A, Fischl B (2003) Regionally localized thinning of the cerebral cortex in Schizophrenia. Archives of General Psychiatry 60:878–888.

Landau WM (2021) The targets R package: a dynamic Make-like function-oriented pipeline toolkit for reproducibility and high-performance computing. Journal of Open Source Software 6:2959.

Lander JP (2018) useful: A Collection of Handy, Useful Functions. In.

Leek JT, Johnson WE, Parker HS, Fertig EJ, Jaffe AE, Zhang Y, Storey JD, Torres LC (2022) sva: Surrogate Variable Analysis. Bioinformatics 17:882–883.

Lerch J (2023) MRIcrotome: Visualization Tools for 3D Volumes. In.

Lerch J, Hammill C, van Eede M, Cassel D (2017) RMINC: Statistical Tools for Medical Imaging NetCDF (MINC) Files. In.

Lerch JP, Sled JG, Henkelman RM (2011) MRI Phenotyping of Genetically Altered Mice. Methods in Molecular Biology:349–361.

Liu S, Seidlitz J, Blumenthal JD, Clasen LS, Raznahan A (2020) Integrative structural, functional, and transcriptomic analyses of sex-biased brain organization in humans. Proc Natl Acad Sci U S A 117:18788–18798.

Mechelli A, Friston KJ, Frackowiak RS, Price CJ (2005) Structural Covariance in the Human Cortex. Journal of Neuroscience 25:8303.

Morris JA, Jordan CL, King ZA, Northcutt KV, Breedlove SM (2008) Sexual dimorphism and steroid responsiveness of the posterodorsal medial amygdala in adult mice. Brain Res 1190:115–121.

Mowinckel AM, Vidal-Piñeiro D (2019) Visualisation of Brain Statistics with R-packages ggseg and ggseg3d. In.

Mowinckel AM, Vidal-Piñeiro D (2022) ggseg: Plotting Tool for Brain Atlases. In.

Mowinckel AM, Vidal-Piñeiro D (2023) ggsegGlasser: Glasser datasets for the ggseg-plotting tool. In.

Neudorfer C, Germann J, Elias GJB, Gramer R, Boutet A, Lozano AM (2020) A high-resolution in vivo magnetic resonance imaging atlas of the human hypothalamic region. Scientific Data 7:305.

Oh SW et al. (2014) A mesoscale connectome of the mouse brain. Nature 508:207–214.

Pedersen TL (2024) patchwork: The Composer of Plots. In.

Persson J, Spreng RN, Turner G, Herlitz A, Morell A, Stening E, Wahlund L-O, Wikström J, Söderlund H (2014) Sex differences in volume and structural covariance of the anterior and posterior hippocampus. NeuroImage 99:215–225.

Pipitone J, Park MTM, Winterburn J, Lett TA, Lerch JP, Pruessner JC, Lepage M, Voineskos AN, Chakravarty MM (2014) Multi-atlas segmentation of the whole hippocampus and subfields using multiple automatically generated templates. NeuroImage 101:494–512.

Qiu LR, Fernandes DJ, Szulc-Lerch KU, Dazai J, Nieman BJ, Turnbull DH, Foster JA, Palmert MR, Lerch JP (2018) Mouse MRI shows brain areas relatively larger in males emerge before those larger in females. Nat Commun 9:2615.

Raznahan A, Lerch JP, Lee N, Greenstein D, Wallace GL, Stockman M, Clasen L, Shaw PW, Giedd JN (2011) Patterns of coordinated anatomical change in human cortical development: a longitudinal neuroimaging study of maturational coupling. Neuron 72:873–884.

Reuter M, Rosas HD, Fischl B (2010) Highly Accurate Inverse Consistent Registration: A Robust Approach. NeuroImage 53:1181–1196.

Richards K, Watson C, Buckley RF, Kurniawan ND, Yang Z, Keller MD, Beare R, Bartlett PF, Egan GF, Galloway GJ, Paxinos G, Petrou S, Reutens DC (2011) Segmentation of the mouse hippocampal formation in magnetic resonance images. NeuroImage 58:732–740.

Robinson D, Hayes A, Couch S (2023) broom: Convert Statistical Objects into Tidy Tibbles. In.

Romero-Garcia R, Whitaker KJ, Váša F, Seidlitz J, Shinn M, Fonagy P, Dolan RJ, Jones PB, Goodyer IM, Bullmore ET, Vértes PE (2018) Structural covariance networks are coupled to expression of genes enriched in supragranular layers of the human cortex. NeuroImage 171:256.

Roos J, Roos M, Schaeffer C, Aron C (1988) Sexual differences in the development of accessory olfactory bulbs in the rat. The Journal of Comparative Neurology 270:121–131.

Rosen AFG, Roalf DR, Ruparel K, Blake J, Seelaus K, Villa LP, Ciric R, Cook PA, Davatzikos C, Elliott MA, Garcia de La Garza A, Gennatas ED, Quarmley M, Schmitt JE, Shinohara RT, Tisdall MD, Craddock RC, Gur RE, Gur RC, Satterthwaite TD (2018) Quantitative assessment of structural image quality. NeuroImage 169:407–418.

Saygin ZM, Kliemann D, Iglesias JE, van der Kouwe AJW, Boyd E, Reuter M, Stevens A, Van Leemput K, McKee A, Frosch MP, Fischl B, Augustinack JC (2017) High-resolution magnetic resonance imaging reveals nuclei of the human amygdala: manual segmentation to automatic atlas. NeuroImage 155:370–382.

Segovia S, Orensanz LM, Valencia A, Guillamón A (1984) Effects of sex steroids on the development of the accessory olfactory bulb in the rat: a volumetric study. Brain Research 16:312–314.

Seitz J, Kubicki M, Jacobs EG, Cherkerzian S, Weiss BK, Papadimitriou G, Mouradian P, Buka S, Goldstein JM, Makris N (2019) Impact of sex and reproductive status on memory circuitry structure and function in early midlife using structural covariance analysis. Human Brain Mapping 40:1221–1233.

Shi Y, Cui D, Niu J, Zhang X, Sun F, Liu H, Dou R, Qiu J, Jiao Q, Cao W, Yu G (2023) Sex differences in structural covariance network based on MRI cortical morphometry: effects on episodic memory. Cereb Cortex 33:8645–8653.

Spencer Noakes TL, Henkelman RM, Nieman BJ (2017) Partitioning k space for cylindrical three dimensional rapid acquisition with relaxation enhancement imaging in the mouse brain. NMR in Biomedicine 30.

Spring S, Lerch JP, Henkelman RM (2007) Sexual dimorphism revealed in the structure of the mouse brain using three-dimensional magnetic resonance imaging. NeuroImage 35:1424–1433.

Steadman PE, Ellegood J, Szulc KU, Turnbull DH, Joyner AL, Henkelman RM, Lerch JP (2014) Genetic Effects on Cerebellar Structure Across Mouse Models of Autism Using a Magnetic Resonance Imaging Atlas. Autism Res 7:124–137.

Ullmann JFP, Watson C, Janke AL, Kurniawan ND, Reutens DC (2013) A segmentation protocol and MRI atlas of the C57BL/6J mouse neocortex. NeuroImage 78:196–203.

Van Essen DC et al. (2012) The Human Connectome Project: A data acquisition perspective. NeuroImage 62:2222–2231.

Vijayakumar N, Ball G, Seal ML, Mundy L, Whittle S, Silk T (2021) The development of structural covariance networks during the transition from childhood to adolescence. Scientific Reports 11:9451.

Wickham H (2020) reshape2: Flexibly Reshape Data: A Reboot of the Reshape Package. In.

Wickham H et al. (2019) Welcome to the tidyverse. Journal of Open Source Software 4:1686.

Wierenga LM, Sexton JA, Laake P, Giedd JN, Tamnes CK (2018) A Key Characteristic of Sex Differences in the Developing Brain: Greater Variability in Brain Structure of Boys than Girls. Cereb Cortex 28:2741–2751.

Wierenga LM et al. (2022) Greater male than female variability in regional brain structure across the lifespan. Human Brain Mapping 43:470–499.

Williams RW, Airey DC, Kulkarni A, Zhou G, Lu L (2001) Genetic Dissection of the Olfactory Bulbs of Mice: QTLs on Four Chromosomes Modulate Bulb Size. Behavior Genetics 31:61–77.

Yang CC, Totzek JF, Lepage M, Lavigne KM (2023) Sex differences in cognition and structural covariance-based morphometric connectivity: evidence from 28,000+ UK Biobank participants. Cereb Cortex 33:10341–10354.

Yee Y, Fernandes DJ, French L, Ellegood J, Cahill LS, Vousden DA, Spencer Noakes L, Scholz J, van Eede MC, Nieman BJ, Sled JG, Lerch JP (2018) Structural covariance of brain region volumes is associated with both structural connectivity and transcriptomic similarity. NeuroImage 179:357–372.

Zaretskaya N, Fischl B, Reuter M, Renvall V, Polimeni JR (2018) Advantages of cortical surface reconstruction using submillimeter 7 T MEMPRAGE. NeuroImage 165:11–26.

