## Supplementary figures and images for "A cross-species analysis of neuroanatomical covariance sex differences in humans and mice"

### Figure 1-1

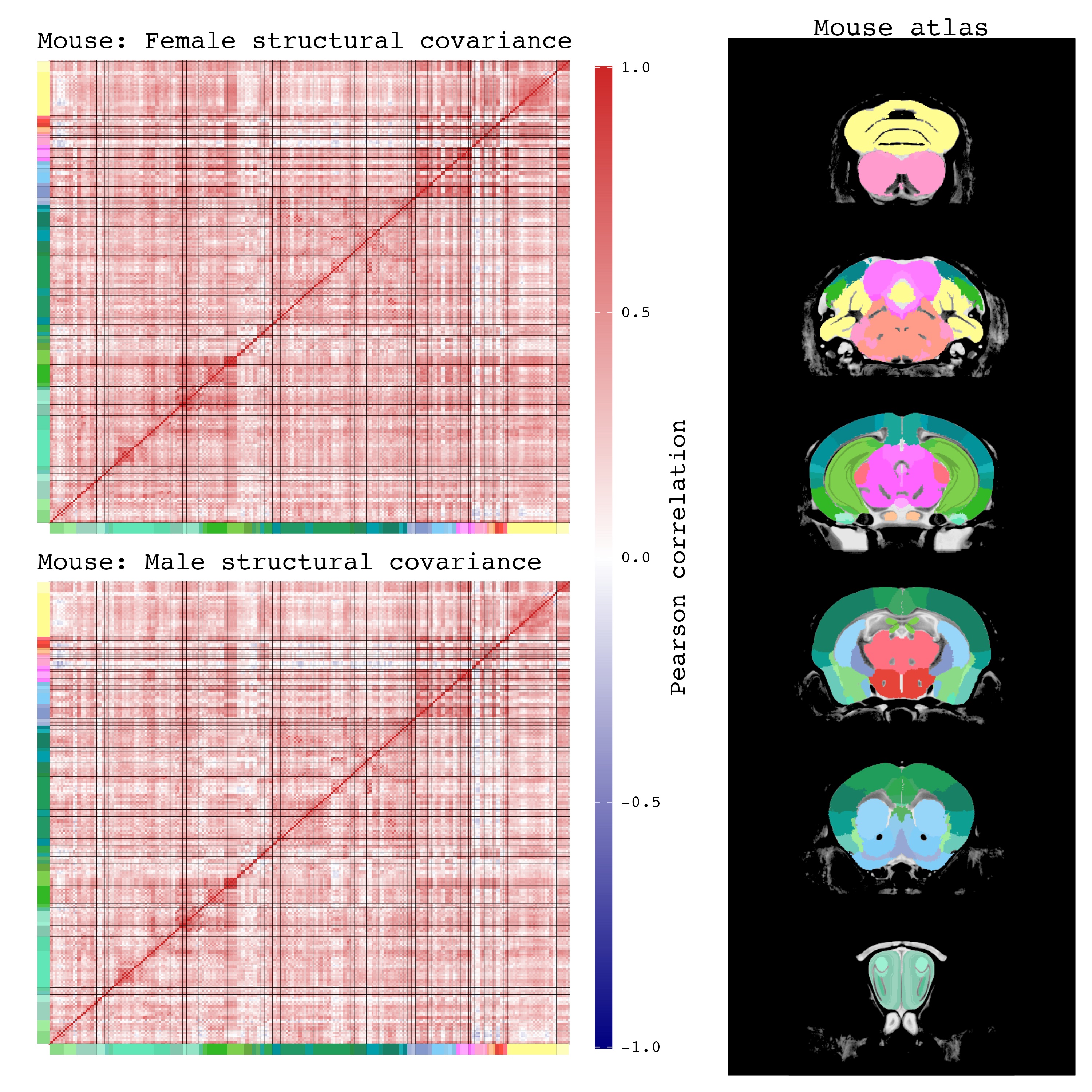

### Figure 1-2

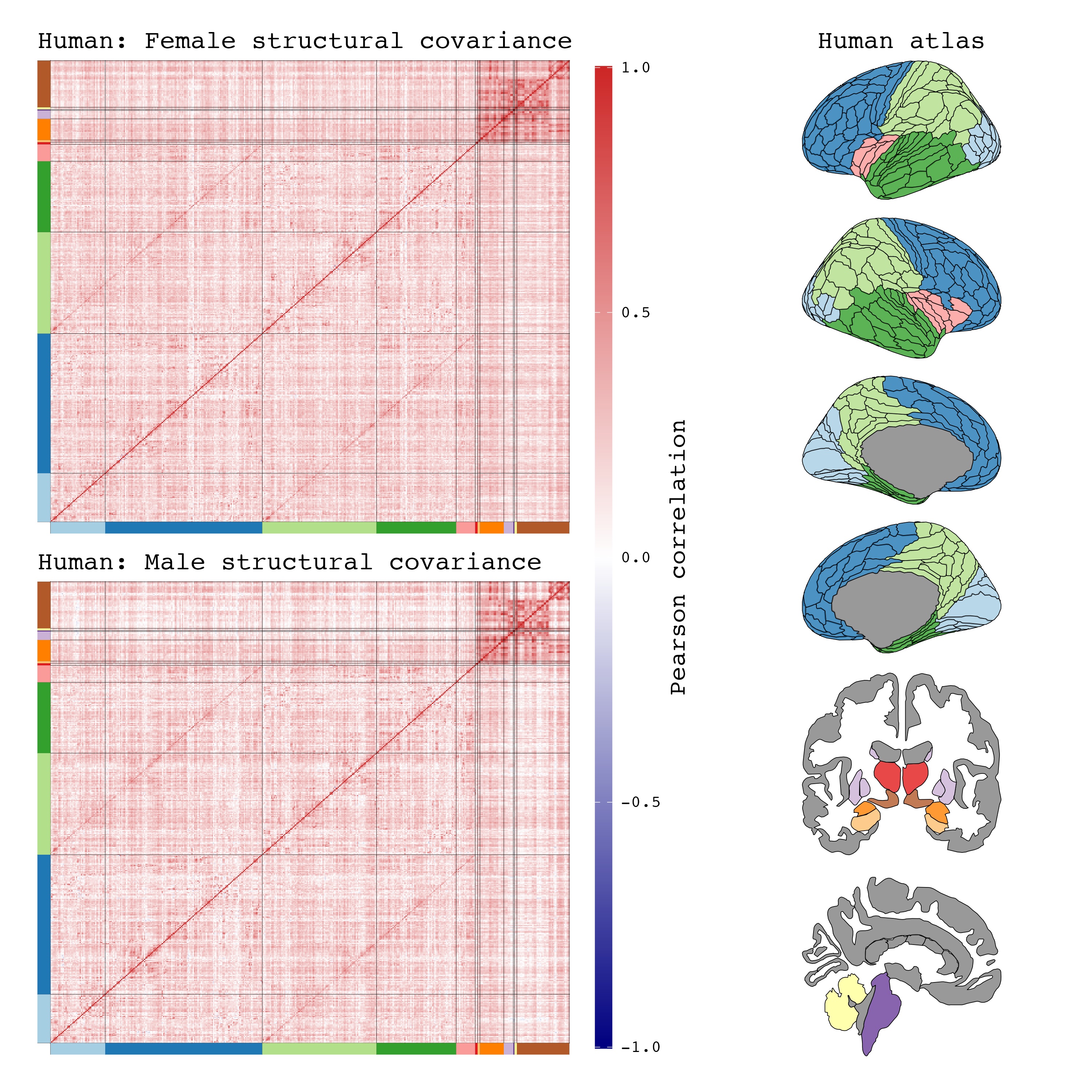

### Figure 1-3

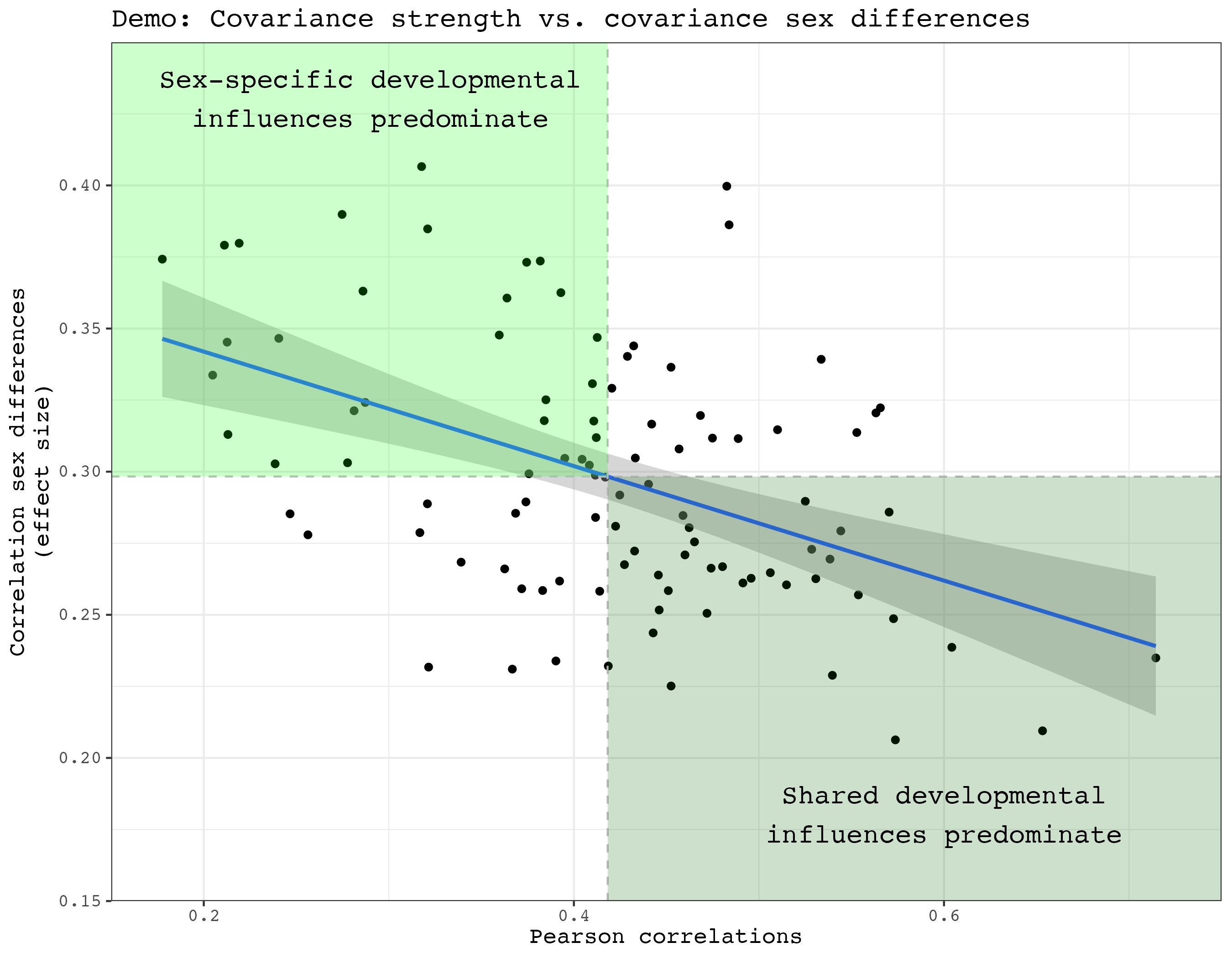

### Figure 1-4

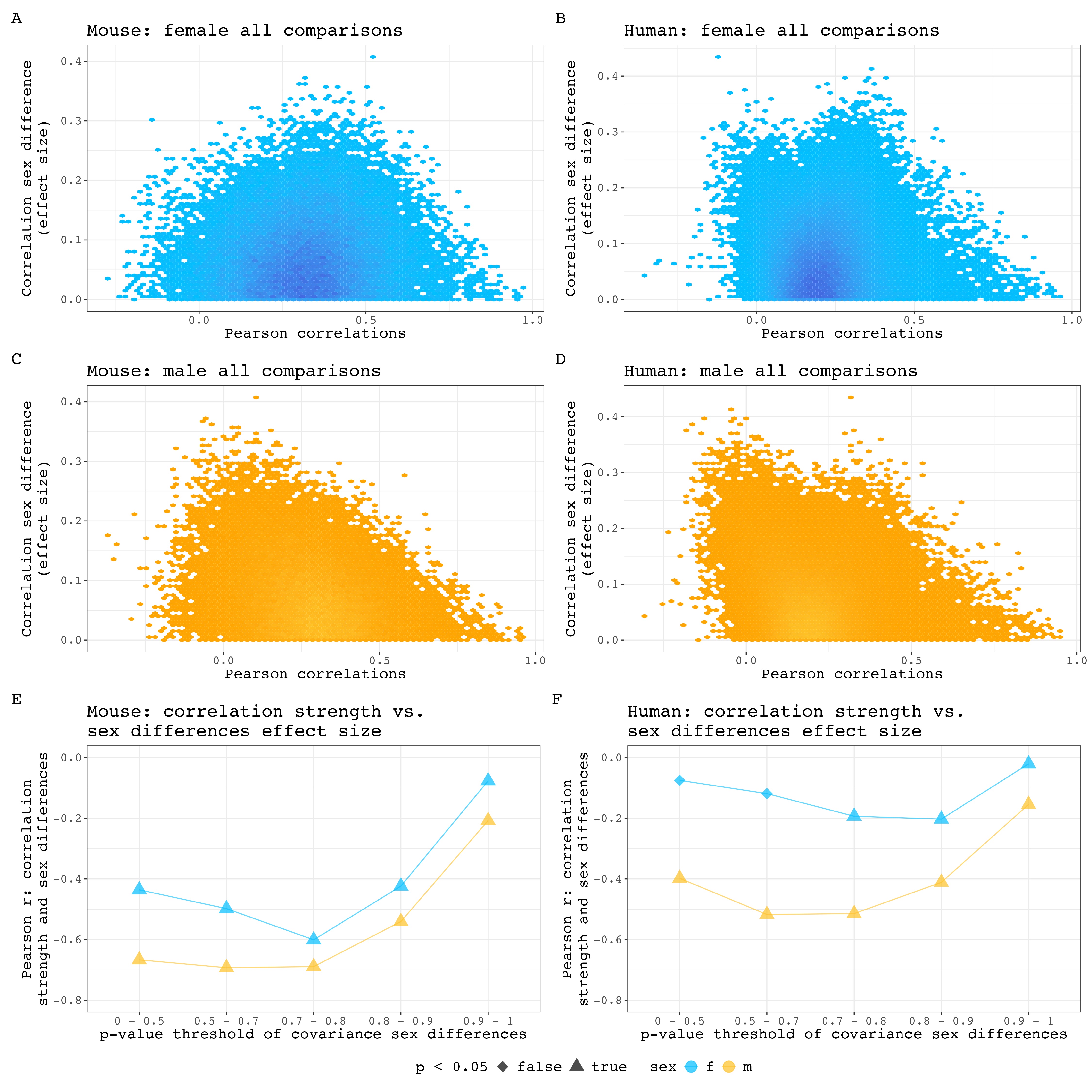
